# Non-enzymatic RNA Glycation is a Metabolic Sensor of Cellular Stress

**DOI:** 10.64898/2026.05.18.725511

**Authors:** Anna Knörlein, Kaila Nishikawa, Naoya Kitamura, Reena Kumari, Shino Murakami, Ankita Shrivastava, Malina Doynova, Nicolae Ciobu, Nicole Walker, Xiaohan Mei, Abdul Vehab Dozic, Jahan Rahman, Yang Xiao, Chiara Evans, Xuejing Yang, Michael G. Kharas, Omar Abdel-Wahab, Francois Fuks, Kamini Singh, Samie Jaffrey, Viraj R. Sanghvi, James J. Galligan, Yael David

**Affiliations:** Chemical Biology Program, Memorial Sloan Kettering Cancer Center, New York, NY, USA; Tri-Institutional PhD Program in Chemical Biology, New York, NY, USA; Department of Pharmacology and Toxicology, College of Pharmacy, University of Arizona, Tucson, AZ, USA; Department of Medicine, Herbert Irving Comprehensive Cancer Center, Columbia University Medical Center, New York, NY, USA; Department of Pharmacology, Weill Cornell Medical College, Cornell University, New York, NY, USA; Department of Molecular Pharmacology, Albert Einstein College of Medicine, Montefiore Einstein Comprehensive Cancer Center, Bronx, NY, USA; Laboratory of Cancer Epigenetics, Faculty of Medicine, ULB-Cancer Research Center (U-CRC), Université libre de Bruxelles (ULB), Institute Jules Bordet, Brussels, Belgium; Computational and Systems Biology Program, Memorial Sloan Kettering Cancer Center, New York, NY, USA; Department of Physiology, Biophysics and Systems Biology, Weill Cornell Medicine, New York, NY, USA; Molecular Pharmacology Program, Sloan Kettering Institute, Memorial Sloan Kettering Cancer Center, New York, NY, USA; Molecular Pharmacology Program, Experimental Therapeutics Center and Center for Stem Cell Biology, Memorial Sloan Kettering Cancer Center, New York, NY, USA

**Keywords:** Metabolism, Methylglyoxal, Glycolysis, RNA modification, Translation, Ribosome Stalling, Ribotoxic Stress Response, Integrated Stress Response

## Abstract

Non-enzymatic RNA modifications expand the epitranscriptome, encoding a rapid and chemistry-driven response to cellular stress. While methylglyoxal, a reactive glycolytic byproduct of metabolic stress, has been shown to modify proteins and DNA, its impact on RNA has remained unexplored. Here, we identify mRNA as a dynamic substrate of MGO, whose modification is actively regulated by DJ-1 and the glyoxalase detoxification system. We show that mRNA glycation impairs translation and engages both the integrated stress response and the ribotoxic stress pathway, culminating in compromised pancreatic β-cell function and reduced insulin secretion. Notably, this phenotype is alleviated by the frontline antihyperglycemic agent metformin. Together, our findings position mRNA as a direct sensor of metabolic stress and establish RNA glycation as a mechanistic link between glycolytic imbalance, translational stress and disease.

## Introduction

Covalent modifications of RNA are central regulators of its structure, stability, and function, shaping nearly every stage of the RNA life cycle.^1^ To date, more than 170 modifications have been identified in both coding and noncoding RNAs, distributed across translated and untranslated regions.^2^ Thus far, most studies have focused on investigating enzymatically installed modifications, which are dynamically written and erased. The most prominent example is *N⁶*-methyladenosine (m⁶A), which plays a key role in regulating mRNA stability and translation.^3,4^ However, alongside these canonical marks, non-enzymatic covalent modifications (NECMs) of RNA have emerged as an additional layer of cellular control.^5^ In contrast to enzymatically installed modifications, NECMs are introduced covalently by reactive molecules generated through cellular metabolism or environmental exposure.^6^ Although predominantly studied in the context of DNA and proteins,^6–8^ increasing evidence suggest that these NECMs on RNA function as sensors of cellular stress, inducing cellular stress pathways and shaping cell fate decisions.^9–11^ For example, ribosomal impairment triggered by UV damage or reactive oxygen species (ROS) was shown to promote eIF2α phosphorylation and activate the Integrated Stress Response (ISR).^9,10^ ISR suppresses cap-dependent translation while selectively promoting the translation of stress-responsive mRNAs, thereby enabling cells to adapt and recover from cellular stress.^11,12^ This ribosomal impairment is also sensed by ZAKα (hereafter referred to as ZAK) which associates to collided ribosomes inducing its auto-phosphorylation and activation.^10,13,14^ Activated ZAK subsequently promotes the phosphorylation and activation of the stress-associated protein kinases p38 and JNK.^9–11^ This so-called ribotoxic stress response (RSR) is triggered by RNA damage caused by UV light and ROS and can lead to cell-cycle arrest, apoptosis, inflammatory responses and metabolic dysregulation.^10,14,15^

Although metabolically driven non-enzymatic glycation has been documented on DNA and proteins, its occurrence and functional consequences on RNA remain largely unexplored. Glycation arises from the non-enzymatic reaction between reducing sugars or metabolic byproducts with proteins and nucleic acids, which is distinguished mechanistically and functionally from enzymatically installed glycosylation.^16^ Among the most potent and reactive endogenous glycating agents is methylglyoxal (MGO), a by-product of glycolysis. MGO is found at low levels (0.5-1μM) in normal cells but is highly elevated in cancer, obesity and plasma of patients with type 2 diabetes mellitus (DM), reaching concentrations of up to 350 μM in disease states.^17–20^ Consistent with this, growing evidence indicates that hyperglycemia-driven accumulation of MGO contributes to the onset and progression of DM by disrupting insulin signaling and secretion, promoting insulin resistance and β-cell dysfunction.^19,21^ Elevated MGO levels and accumulation of glycation are linked to DM complications including endothelial dysfunction, inflammation, and aberrant angiogenesis, pathologies that drive vascular injury and cardiovascular complications, the leading cause of mortality in patients with DM.^22^ Intracellular MGO is primarily detoxified by the glyoxalase system, composed of GLO1 and GLO2, which converts MGO into lactate. However, this detoxification pathway is frequently downregulated in patients with DM.^19,23,24^ Moreover, it was shown that the Parkinson’s-associated protein DJ-1 can reduce glycation levels with indications to both glyoxalase and deglycase activities on proteins, nucleotides, and nucleic acids.^25,26^ MGO can additionally be scavenged by short regulatory peptides, such as glutathione, carnosine, as well as by small molecules.^19,27,28^ Although glycation of proteins and DNA is established as a driver of cellular dysfunction, genomic instability, epigenetic alterations, and malignant transformation,^29–31^ little is known about RNA glycation. It is unclear how prevalent RNA glycation is as a cellular modification, what is its biological significance and contribution to cellular homeostasis and disease pathogenesis. Similarly to DNA, MGO modifies the exocyclic amine of guanosine leading to multiple chemically distinct RNA glycation adducts. The two major products are N²-carboxyethyl-2′-guanosine (CEG) and the cyclic adduct C-MGO-G, which together are collectively termed G_MGO_ (Fig. 1A) and serve in this study as a representative readout of RNA glycation.^28,32^ Despite limited mechanistic insight, free, extracellular CEG was detected in both serum and urine of patients with DM or pre-diabetes, suggesting its potential as a biomarker for disease state.^33,34^ Moreover, patients who developed DM complications exhibited even higher levels of free CEG in their urine,^33^ supporting a correlation between disease progression and RNA glycation *in vivo*. Finally, preliminary evidence shows that RNA glycation decreases translation and RNA stability.^33,35^ However, these studies measure systemic levels rather than RNA glycation events occurring directly within cells, leaving the intracellular formation and regulation of RNA glycation, as well as its broader molecular consequences and causal role in disease progression, largely undefined.

**Fig. 1:**
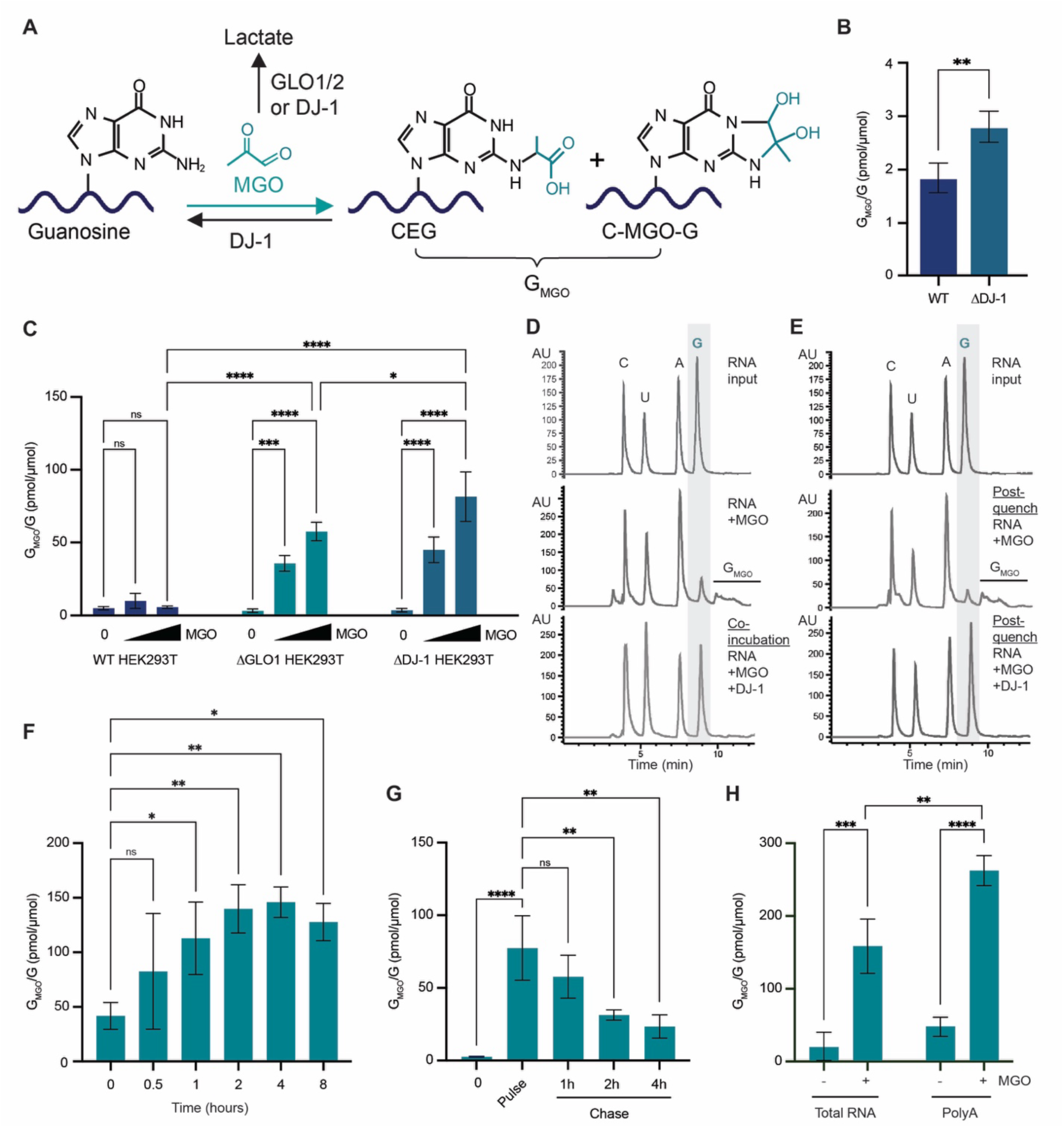
RNA glycation accumulates and is regulated *in vitro* and *in vivo*. (A) Schematic illustrating the proposed regulation of RNA glycation by methylglyoxal (MGO), GLO1/2 and DJ-1. (B) Levels of MGO derived guanosine adducts (G_MGO_) in total RNA isolated from liver tissue of wild-type (WT) and DJ-1 deficient (ΔDJ-1) mice, quantified by LCMS relative to isotopically labeled G or G_MGO_ internal standards (n=5). Statistical significance was assessed by two-tailed Welch’s *t* test. (\**P*< 0.0332; \*\**P*< 0.0021). (C) G_MGO_ levels of total RNA isolated from WT, glyoxalase-deficient (ΔGLO1), and DJ-1-deficient (ΔDJ-1) HEK293T cells treated with 0, 24 µM, 60 µM MGO for 1 hour. Quantified by LCMS to isotopically labeled standards (n=3). Statistical significance was assessed using a two-way ANOVA with Tukey’s multiple comparison test (\**P*< 0.0332; \*\**P*< 0.0021; \*\*\**P*< 0.0002; \*\*\*\**P*< 0.0001). (D) RP-HPLC chromatogram of digested nucleotides from 5 µg total RNA, 5 µg RNA incubated with 1.2 mM MGO for 16 hours, and 5 µg RNA co-incubated with 1.2 mM MGO and 4 µM DJ-1for 16 hours at 37 °C (AU, arbitrary units) (n=3). (E) RP-HPLC chromatogram of digested nucleotides from 5 µg total RNA, 5 µg RNA preglycated with 1.2 mM MGO for 16 hours then quenched with 25 mM Tris, and 5 µg RNA preglycated with 1.2 mM MGO for 16 hours, quenched with 25 mM Tris, then incubated with 4 µM DJ-1 for 16 hours at 37 °C (AU, arbitrary units) (n=3). (F) G_MGO_ levels of total RNA isolated from ΔGLO1 HEK293T cells treated with 60 µM MGO for the indicated durations. Quantified by LCMS (n=3). Statistical significance was assessed using a one-way ANOVA with Dunnett’s multiple comparison test (\**P*< 0.0332; \*\**P*< 0.0021; \*\*\**P*< 0.0002; \*\*\*\**P*< 0.0001). (G) G_MGO_ levels of total RNA isolated ΔGLO1 HEK293T cells subjected to an MGO pulse–chase assay (1 hour pulse with 60 µM MGO followed by media change and measured after the indicated times) quantified by LCMS (n=3). Statistical significance was assessed using a Two-way ANOVA with Tukey’s multiple comparison test (\**P*< 0.0332; \*\**P*< 0.0021; \*\*\**P*< 0.0002; \*\*\*\**P*< 0.0001). (H) G_MGO_ levels of total and polyA-enriched RNA isolated from ΔGLO1 HEK293T cells treated with 0 and 60 µM MGO for 4 hours. Quantified by LCMS (n=3). Statistical significance was assessed using a Two-way ANOVA with Tukey’s multiple comparison test (\**P*< 0.0332; \*\**P*< 0.0021; \*\*\**P*< 0.0002; \*\*\*\**P*< 0.0001). All data are presented as mean ± SD.

Here, we perform a comprehensive fundamental study that led to the identification of RNA glycation as a metabolically responsive modification that dynamically links glycolytic flux to translational control and acts as a sensor of key cellular functions in DM. Using a highly interdisciplinary approach including quantitative mass spectrometry, *in vitro* biochemical assays, and cellular and mouse models, we determine the susceptibility of certain mRNAs to glycation upon metabolic stress. These RNA glycation events trigger both ISR and RSR through ribosomal impairment. Together, these results offer a new mechanism by which RNA glycation impairs insulin secretion via the activation of RSR and contributes to DM progression and can be directly rescued by the hyperglycemic drug metformin. Importantly, our findings reveal that RNA glycation represents a previously unrecognized link between cellular metabolic state to translational control, providing a unifying framework for several previously disconnected observations.

## Results

### RNA Glycation is a dynamically regulated modification *in vitro* and *in vivo*

While CEG has been detected as free, extracellular nucleotides in the serum and urine of diabetic patients,^33,34^ how glycation accumulates and is regulated on intracellular RNA remains largely unexplored. To address this gap, we first synthesized isotopically labeled standards of unmodified G and G_MGO_ to allow for precise quantification of intracellular RNA glycation by liquid chromatography-mass spectrometry (LCMS) (Fig. S1A). Using this approach, we determine RNA glycation levels *in vivo* using liver tissue from wild-type (WT) and DJ-1 knockout (ΔDJ-1) mice. Liver tissue was selected for analysis due to its endogenously elevated MGO levels and central role in glycolysis and MGO detoxification, making it a site of heightened metabolic stress.^36^ DJ-1 knockout liver tissue was chosen due to the central role it plays in MGO clearance and disease.^25,37^ Indeed, LCMS analysis revealed a substantial increase in intracellular G_MGO_ levels in ΔDJ-1 liver total RNA samples compared to WT controls (Fig. 1B), demonstrating that DJ-1 limits RNA glycation under physiological conditions (Fig. 1A).

Because *in vivo* analyses largely capture steady-state accumulation and thus would not resolve RNA glycation kinetics or short-term regulation, we next used a similar methodology to evaluate glycation levels in cultured cells. To assess the contribution of the canonical endogenous detoxification systems to regulating RNA glycation, we analyzed RNA extracted from cultured HEK293T cells lacking glyoxalase enzyme GLO1 (ΔGLO1) or DJ-1 (ΔDJ-1).^38^ Cells were exposed to MGO at levels measured in disease states^20^ after which total RNA was extracted and analyzed by LCMS. Our results indicate elevated levels of G_MGO_ in both deletion strains compared with WT (Fig. 1C). Notably, WT HEK293T cells reach similar accumulation of RNA glycation but at higher MGO concentrations (Fig. S1H) or longer exposure times (4 h, Fig. S1I), indicating that DJ-1- and GLO1-mediated detoxification pathways substantially attenuate, but do not fully eliminate, RNA glycation in normal cells.

To determine whether DJ-1 functions as a glyoxalase that quenches MGO prior to its reaction with RNA or can directly decrease the levels of RNA glycation,^39^ we analyzed dynamics of RNA glycation in the absence or presence of recombinant DJ-1. Briefly, total RNA was isolated from WT HEK293T cells, then incubated with MGO alone or co-incubated with recombinant DJ-1,^29^ followed by digestion to mononucleotides and HPLC analysis. Our results show that MGO treatment induced the accumulation of G_MGO_ as indicated by a selective reduction in the guanosine peak accompanied by the emergence of multiple new peaks corresponding to glycation adducts on guanosine (Fig. 1D). In contrast, co-incubation of RNA with MGO and recombinant DJ-1 yielded a chromatogram lacking glycation peaks, mirroring that of unmodified RNA, indicating that DJ-1 prevents the formation of RNA glycation adducts by MGO *in vitro* (Fig. 1D). Importantly, treating pre-glycated, purified RNA with recombinant DJ-1 resulted in a similarly reduced RNA glycation to levels comparable to untreated RNA, indicating that DJ-1 can also directly impact RNA glycation adducts post-formation *in vitro* (Fig. 1E).

We next sought to investigate the dynamic regulation of glycated RNA in cells. To enhance glycation signal detection, we utilized ΔGLO1 HEK293T cells, selected over ΔDJ-1 in order to minimize confounding effects associated with DJ-1’s multiple cellular functions.^37,39,40^ We first performed a time-course experiment in which cells were treated with MGO and analyzed at the indicated times. Our results show that G_MGO_ rapidly accumulates within 1 hour and peaks after approximately 4 hours following MGO exposure. RNA glycation begins to decline at 8 hours, consistent with RNA turnover (Fig. 1F). Notably, even in the absence of exogenous MGO, we detected basal levels of RNA glycation in ΔGLO1 HEK293T cells, suggesting that physiological MGO production is sufficient to modify RNA when detoxification pathways are impaired. To determine the stability of G_MGO_, we performed a pulse-chase experiment. Cells were exposed to MGO for 1 hour, then media was changed to MGO-free media, and G_MGO_ levels were monitored over time. Our results indicate that G_MGO_ levels decline substantially within 2 hours following an MGO pulse, indicating that RNA glycation is highly dynamic and maintained by a balance between glycation, RNA turnover, and active removal of glycation adducts (Fig. 1G).

Having established its accumulation on RNA, we next investigated the subcellular distribution and RNA species that are most susceptible to glycation. Cellular fractionation revealed comparable glycation levels in nuclear and cytoplasmic RNA (Fig. S1J), supporting RNA glycation occurs broadly rather than restricted to a specific compartment. To identify the specific RNA species that undergo glycation, we treated cells with MGO and enriched for polyadenylated RNA.^41^ Comparison of glycation levels between poly(A)-enriched RNA and total RNA revealed higher glycation in the mRNA-enriched fraction (Fig. 1H). Because mRNA constitutes only a small fraction of total cellular RNA compared to rRNA and tRNA, these results indicate a selective and disproportionate accumulation of glycation within mRNA. Collectively, our results establish RNA glycation as a metabolically responsive RNA modification. Similarly to DNA and protein glycation, RNA glycation is regulated by the glyoxalase pathway and the dual protective–repair activities of DJ-1. However, in contrast to the slow accumulation of protein adducts, RNA glycation rapidly appears, particularly on mature mRNA transcripts, and quickly turns over, suggesting that it functions at short time scales, linking enhanced glycolytic flux to dynamic regulation at the RNA level.

### Glycation primarily accumulates in coding regions of mRNA

After establishing that mature mRNA is the major RNA species that accumulates glycation, we next sought to determine the precise transcripts and sites of mRNA glycation. To do so, we leveraged a published dataset which employed a kethoxal probe to map modified guanosines by enriching for kethoxal-reactive sites followed by RNA-sequencing (Keth-seq).^42^ Since kethoxal is chemically highly similar to MGO and reacts with guanosine residues with similar chemistry, it serves as a proxy for nucleotides susceptible to glycation (Fig. 2A). Analysis of the Keth-seq datasets generated from HeLa cells revealed that most guanosines exhibit low reactivity (Fig. 2B). However, a distinct subset of transcripts display high reactivity scores (defined by a score ≥ 0.5), are therefore more frequently or more efficiently modified, consistent with selective targeting of specific sites.^42,43^ Further analysis of reactive guanosines confirmed that the number of reactive guanosines per transcript correlates positively with transcript length (R = 0.51, p < 2.2 × 10⁻¹⁶; Fig. 2C). Importantly, only a small number of transcripts contained high levels (more than 30%) of reactive guanosines (Fig. 2D), but these reactive guanosines were mostly enriched within coding sequences (47.1% Fig. 2E). As an example, we plotted the kethoxal reactivity scores against the position in the transcript of one of the highly modified transcripts, NOP10 (Fig. 2F). Except for two adjacent reactive non-G positions, likely due to mapping artifacts, only guanosines were reactive, consistent with the high specificity of kethoxal towards guanosine. Within NOP10 only a subset of guanosines was reactive, demonstrating a selective modification rather than general reactivity for guanosines, and those are located primarily in the CDS and 3’ UTR regions.

**Fig. 2:**
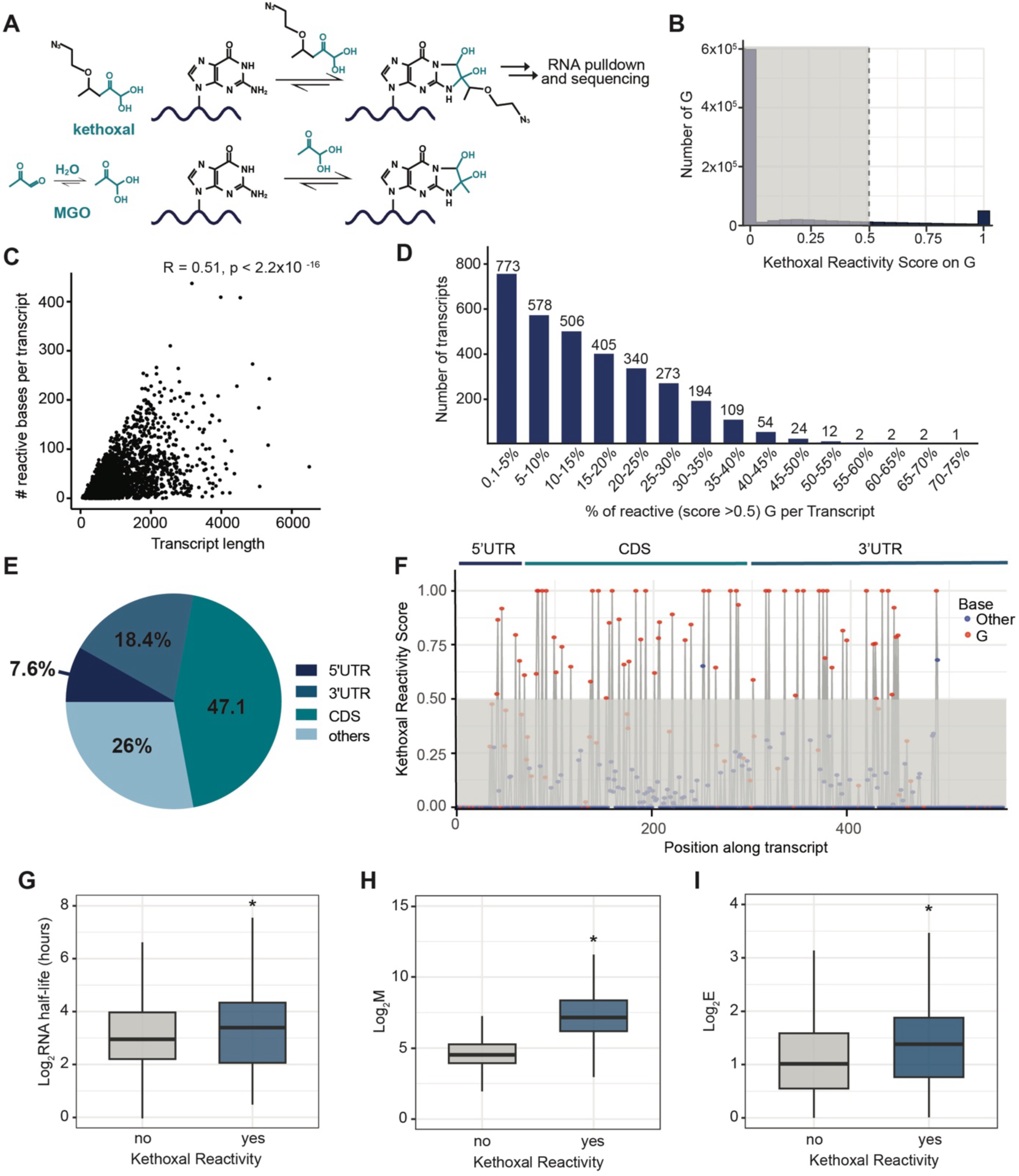
A distinct subset of RNAs is highly susceptive to glycation and accumulates on coding regions. (A) Schematic of Keth-seq workflow involving the reaction of N_3_-kethoxal and guanine.^42^ MGO reaction with guanine shown for structural and reactivity comparison. (B) Reanalysis of kethoxal reactivity scores for all guanosine residues detected transcriptome-wide in HeLa cells.^42^ Guanosines with a reactivity score > 0.5 were classified as reactive (dashed line). (C) Relationship between transcript length and the number of kethoxal-reactive guanosines per transcript (reactivity score > 0.5). Each dot represents one transcript; Pearson correlation coefficient and *P* value are indicated. (D) Distribution of transcripts based on the percentage of reactive guanosines (> 0.5) per transcript. Bars indicate the number of transcripts falling within each reactivity bin. (E) Transcriptome-wide localization of kethoxal-reactive guanosines. Pie chart shows the percentage of reactive sites (> 0.5) within 5′ UTR, CDS, 3′ UTR, and other non-annotated regions. (F) Nucleotide-resolution kethoxal reactivity profile across the *NOP10* transcript. Reactivity scores for individual bases are shown along transcript coordinates (with bases >0.5 classified as reactive (dashed line)) with annotation of the estimated 5′ UTR, CDS, and 3′ UTR regions. (G) Comparison of RNA half-life (hours) between kethoxal-reactive (at least one G>0.5) and non-reactive transcripts. Box plots show median, interquartile range, and whiskers; A Kolmogorov–Smirnov test was performed to evaluate differences in the distribution (H) Comparison of RNA abundance (M) between kethoxal-reactive (at least one G>0.5) and non-reactive transcripts. Box plots show median, interquartile range, and whiskers; A Wilcoxson sum ranked test was performed to evaluate differences in the distribution, with *P*<0.05. (I) Comparison of RNA translation efficiency (E) between kethoxal-reactive (at least one G>0.5) and non-reactive transcripts. Box plots show median, interquartile range, and whiskers; A Wilcoxson sum ranked test was performed to evaluate differences in the distribution, with *P*<0.05.

Notably, transcripts harboring reactive guanosines exhibited significantly increased RNA half-lives (Fig. 2G) and overall abundance (Fig. 2H) compared to nonreactive species (p < 0.05). These results suggest a link between transcript stability and RNA levels and the accumulation of RNA glycation, which is characteristic of non-enzymatic modifications.^5,44^ In addition, we observed a correlation between highly reactive transcripts and increased translation efficiency (Fig. 2I) indicating that glycation impacts highly translated mRNAs more than lowly translated ones. Although translation efficiency is shaped by multiple parameters,^45^ this association further suggests that mRNAs that possess intrinsic properties that favor translation such as reduced secondary structure and increased stability, are more susceptible to glycation. Together, these data indicate that RNA glycation accumulates on a subset of transcripts and guanosine residues rather than occurring stochastically, with patterns that reflect RNA length, abundance, structure and stability.^42^ Glycation-prone guanosines are enriched within coding sequences and preferentially occur in long-lived and abundant mRNAs, consistent with increased exposure to reactive metabolites.

### mRNA glycation stalls translation *in vitro* and *in cellulo*

Given that RNA glycation accumulates rapidly on highly translated mRNAs, we next asked whether this modification has functional consequences for protein synthesis. Although preliminary studies showed that mRNA glycation suppresses translation *in vitro*,^33,35,46^ we sought to perform a detailed mechanistic investigation of the direct effects of RNA glycation on translation. First, we determined the global impact of short-term exposure of MGO on cellular translation. We treated both WT and ΔGLO1 HEK293T cells with increasing concentrations of MGO and monitored translation using homopropargylglycine (HPG), a noncanonical methionine analog that marks newly synthesized proteins. In WT cells, MGO induced a dose-dependent translation arrest reaching levels comparable to those observed following treatment with the protein synthesis inhibitor anisomycin (ANS)^47^ (Fig. 3A). ΔGLO1 cells exhibit a lower basal level of translation consistent with elevated endogenous glycation levels and had complete inhibition of translation upon MGO treatment (Fig. 3A). To further validate the impact of glycation on global translation, we utilized a puromycin incorporation assay, where puromycin is incorporated into nascent polypeptide chains and can subsequently detected by immunoblot. Indeed, treatment with increasing concentrations of MGO led to a robust inhibition of bulk translation in both WT and ΔGLO1 cells, comparable to anisomycin, as evidenced by a decrease in puromycin incorporation (Fig. 3B). We did not detect reduced basal translation in ΔGLO1 cells compared to WT cells, possibly due to lower assay sensitivity. Notably, we did not detect protein glycation by Western blot under these experimental conditions (Fig. S4C) suggesting that the effects on cellular translation are specifically induced by RNA and not protein glycation. Using the puromycin assay, we also demonstrate that the impact of glycation on global translation is reversible, with protein synthesis resuming 2–4 hours after a chase with MGO-free media (Fig. S2A), consistent with the decrease in RNA glycation observed in Fig. 1G.

**Fig. 3:**
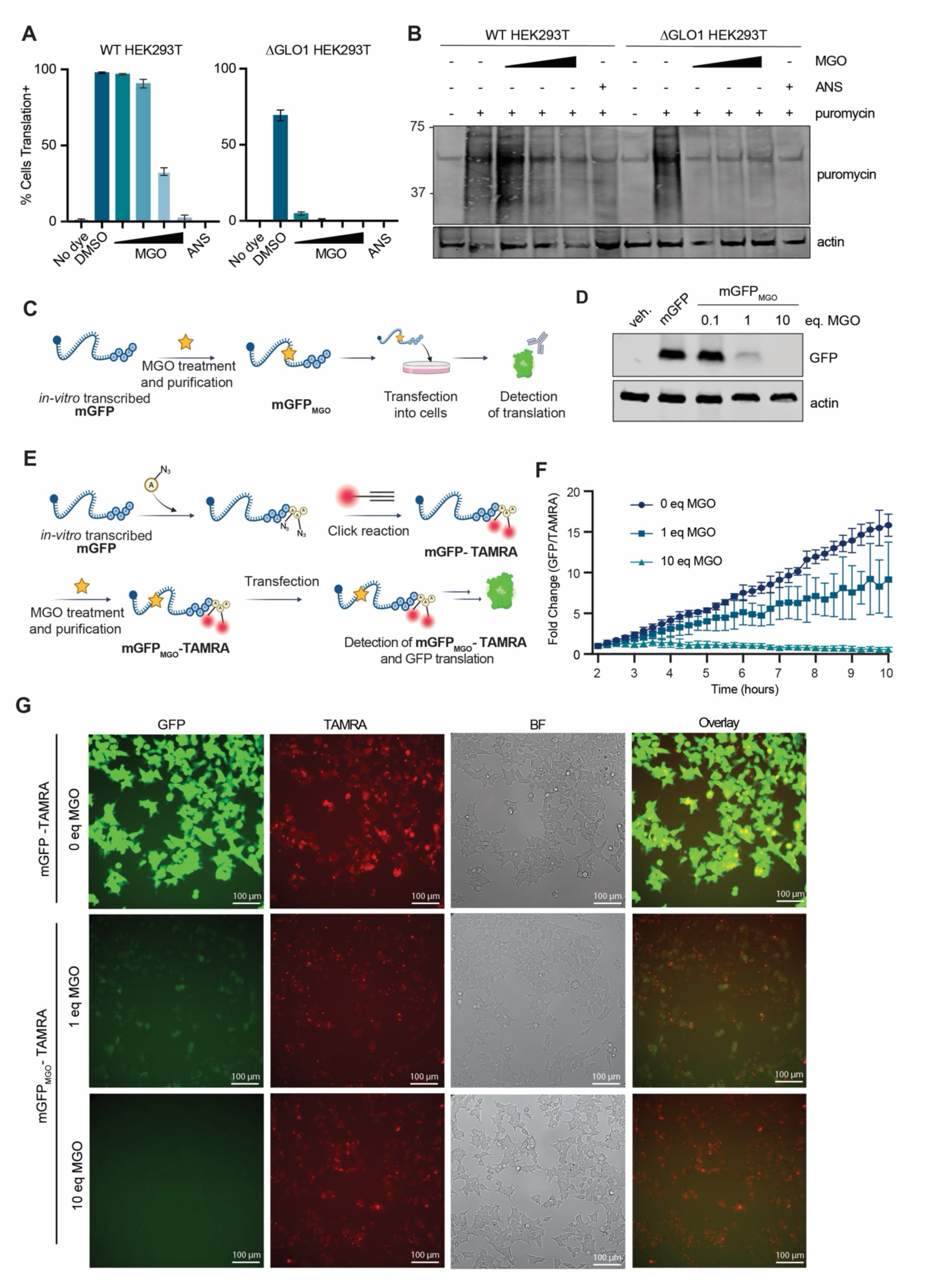
mRNA glycation suppresses translation. (A) HPG incorporation assay of percent translation rate in WT and ΔGLO1 HEK293T cells treated with 0-0.47 mM MGO (WT) and 0-0.12 mM MGO (ΔGLO1) (n=3). Anisomycin (1 µg/mL) was used as a control for translation inhibition. (B) Immunoblot of puromycin incorporation in WT and ΔGLO1 HEK293T cells. Cells were treated with 0-0.47 mM MGO (WT) and 0-0.12 mM MGO (ΔGLO1) or 1 µg/mL anisomycin for 45 min before media was exchanged to media containing 5 µg/mL puromycin for 15 min (n=3). (C) Schematic of mGFP and mGFP_MGO_ preglycation, transfection, and GFP detection. (D) Immunoblot of WT HEK293T cells transfected for 4 hours with a vehicle control, mGFP or mGFP_MGO_ (0-10 equivalents of MGO per guanosine for 16 hours at 37 °C) (n=3). (E) Schematic of mGFP labelling with TAMRA (mGFP-TAMRA) and preglycation (mGFP_MGO_-TAMRA) for dual detection during live cell imaging. (F) Quantification of live-cell imaging of WT HEK293T cells 10 hours after transfection with either TAMRA–mGFP or TAMRA–mGFP_MGO_ (modified with 1 or 10 equivalents of MGO per guanosine for 16 h at 37 °C). GFP fluorescence was quantified and normalized to the TAMRA signal, and data are presented as fold change (Data represents mean ± SD from 4 technical replicates). (E) Representative live-cell images of WT HEK293T cells 10 hours post transfection with either TAMRA-mGFP or TAMRA-mGFP_MGO_ (1 or 10 equivalents of MGO per guanosine for 16 hours at 37 °C) (n=3).

Although we detect RNA glycation coincident with translational inhibition, preceding any detectable accumulation of glycated proteins, we sought to determine whether the observed suppression of global translation by MGO is directly linked mRNA glycation. For that, we synthesized a GFP-encoding mRNA reporter (mGFP). Briefly, the GFP mRNA transcript was generated by *in vitro* transcription using a linearized DNA template followed by *in vitro* capping to generate a functional mRNA. The mature mGFP was incubated with MGO at either 1 equivalent (sub-stoichiometric conditions) or 10 equivalents (excess MGO) and purified to generate glycated mRNA (mGFP_MGO_). Either unglycated mGFP or glycated mGFP_MGO_ was transfected into WT HEK293T cells, where GFP translation was detected by Western blot analysis (Fig. 3C). Indeed, we observed a dose-dependent decrease in GFP protein levels in cells corresponding to increasing levels of mGFP_MGO_ preglycation, indicating reduced translation of mGFP_MGO_ (Fig. 3D). To control for potential differences in transfection efficiency arising from mRNA glycation, we fluorescently labeled the 3′ poly(A) tail of the GFP-encoding mRNA, a region not susceptible to glycation, prior to MGO treatment (mGFP-TAMRA). This strategy enables direct monitoring of RNA delivery independent of glycation status (Fig. 3E). WT HEK293T cells were transfected with mGFP-TAMRA or mGFP_MGO_-TAMRA and monitored for RNA uptake (TAMRA) and translational output (GFP) by live cell imaging. Normalizing the GFP signal to the reduced transfection efficiency of glycated mRNA revealed a quantitative decrease in GFP translation, which correlated with the extent of mGFP preglycation (Fig. 3F, G). Altogether these analyses demonstrate that RNA glycation suppresses protein synthesis in cells, establishing it as a direct, physiologically relevant mechanism for modulating translation under cellular stress.

### mRNA glycation induces ribosome collisions and the integrated stress response (ISR)

We next sought to determine the underlying mechanism by which MGO impairs protein synthesis. We hypothesized that glycation, like other RNA NECMs, acts as a “roadblock” for the translation machinery, leading to ribosomal impairment and stalling of translation.^9,48,49^ To investigate whether elevated levels of MGO induce ribosome impairment, we first followed RPS3 monoubiquitination, which accumulates upon ribosome collision on cytosolic mRNAs.^50,51^ Western blot analysis of WT HEK293T cells treated with MGO for 1 hour revealed the appearance of a second, higher-molecular-weight band consistent with monoubiquitinated RPS3 (Fig. 4A).^50,51^ Next, we examined whether this impairment is a direct consequence of RNA glycation. We performed *in vitro* ribosome collision assays using [³²P]-labeled unglycated mGFP and glycated mGFP_MGO_ in HeLa cell lysate.^52^ The ribosome footprint analysis revealed a concentration-dependent increase in disomes and trisomes formation with increasing levels of mRNA glycation (Fig. 4B) suggesting that RNA glycation is sufficient to impair elongation and promote ribosome collisions.

**Fig. 4:**
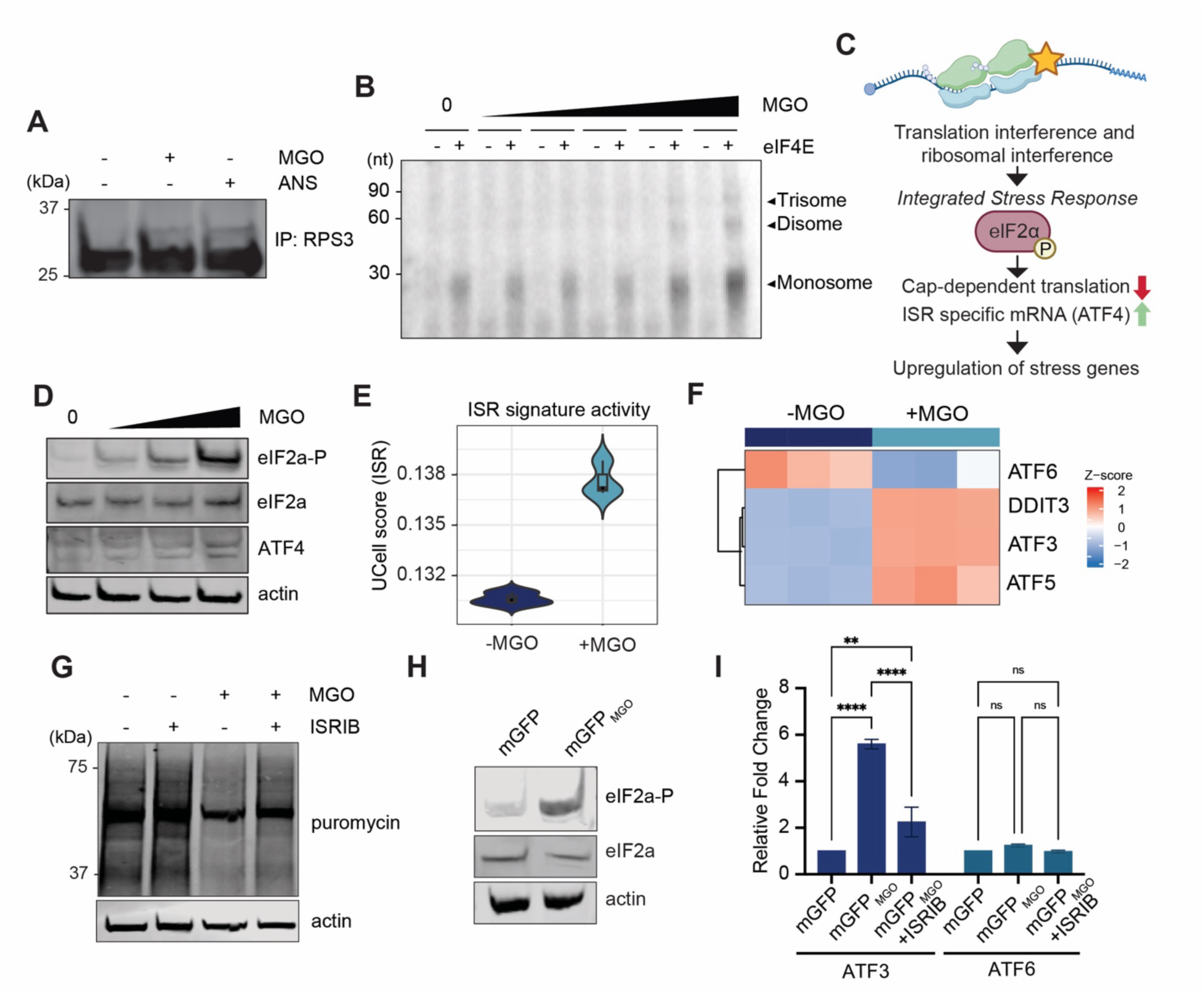
RNA glycation induces ribosome collision and triggers the integrated stress response. (A) Immunoprecipitation of ribosomal protein RPS3 followed by immunoblotting for RPS3 from WT HEK293T cells treated with 0 or 0.24 mM MGO for 1 h or 1 µg/mL anisomycin for 1 h demonstrating ribosome collision–associated RPS3 ubiquitination (n=3). (B) *In vitro* translation assay using GFP mRNA subjected to increasing levels of glycation. *In vitro*–transcribed GFP mRNA was incubated with increasing levels of MGO (0–2.35mM) for 6 hours, purified, and translated in HeLa cell lysate. Polysome distributions reveal dose-dependent accumulation of stalled ribosomes. (C) Schematic illustrating the activation of integrated stress response (ISR) in response to RNA glycation–induced ribosome stalling. (D) Immunoblots showing eIF2α phosphorylation and ATF4 upregulation after incubation of WT HEK293T cells for 1 hour with 0-0.24 mM MGO (n=3). (E) ISR activation signature derived from RNA-seq data, comparing control and MGO-treated ΔGLO1 HEK293T cells (60 µM MGO, 4 h). Each violin represents the distribution of ISR signature scores across samples. (F) Gene expression changes associated with ISR activation in ΔGLO1 HEK293T cells treated with 60 µM MGO for 4 hours, as determined by RNA seq. Heatmap shows Z-score–normalized expression of canonical ISR and stress-responsive genes. (G) Immunoblot of puromycin incorporation in WT HEK293T cells. Cells were pretreated with 200 nM ISRIB for 30 min, followed by treatment with MGO 0 or 0.12mM) in the presence of ISRIB for an additional 45 min. Media was then replaced with puromycin-containing media (5 µg/mL) for 15 min to label nascent polypeptides (n=3). (H) Immunoblot analysis of WT HEK293T cells transfected for 4 hours with either mGFP or mGFP_MGO_ (10 equivalents of MGO per guanosine for 16 hours at 37 °C) (n=3). (I) Relative mRNA expression of ATF3 and ATF6 in WT HEK293T transfected for 4 hours with either mGFP or mGFP_MGO_ (10 equivalents of MGO per guanosine for 16 hours at 37 °C) in the presence or absence of ISRIB, as indicated. Transcript levels were quantified by qPCR and normalized to housekeeping gene expression, then expressed relative to the untreated control. Data represent *n* = 2 biological replicates with 3 technical replicates each and are shown as mean ± SD. Statistical significance was assessed using two-way ANOVA with Tukey’s multiple-comparison test (*P* < 0.0332; \**P* < 0.0021; \*\**P* < 0.0002; \*\*\**P* < 0.0001).

Ribosome impairment is detected by several cellular surveillance pathways that initiate stress-adaptive signaling programs including the ISR. The hallmark of ISR is phosphorylation of eIF2α at Ser51, leading to global suppression of cap-dependent translation while selectively enhancing translation of stress-responsive transcripts such as ATF4. ATF4 orchestrates transcriptional programs that promote amino acid biosynthesis, antioxidant defenses, autophagy, and protein quality control, thereby enabling cells to adapt and recover from divers stressors including ribosomal perturbations, nutrient deprivation and oxidative stress (Fig. 4C).^53^ To test whether elevated MGO levels induce activation of the ISR, we treated WT HEK293T cells with increasing concentrations of MGO for 1 hour followed by Western blot analysis of eIF2α and ATF4. Our results indicate an MGO dose-dependent activation of eIF2α as well as increased levels of ATF4 (Fig. 4D), consistent with downstream ISR activation. We next asked whether elevated MGO levels also elicits ISR-dependent changes to the cellular transcriptional programming. To address this, we performed a global RNA-seq following MGO treatment. To maximize sensitivity for detecting MGO-dependent changes, we used ΔGLO1 HEK293T cells and 4 hours of MGO treatment, which showed the highest G_MGO_ accumulation in our assays (Fig. 1F). Our results indicate that in total, about 60% of ISR-associated genes were upregulated following MGO treatment, demonstrating a substantial translational reprogramming upon ISR activation (Fig. 4E, Fig. S3). Specifically, we observed an upregulation of key ISR-associated genes ATF3, ATF5, and DDIT3 (Fig. 4F). In contrast, the expression of ATF6, a transcription factor activated during the ER unfolded protein response, was not increased suggesting that the observed ISR activation occurs independently of ATF6-mediated ER stress signaling (Fig. 4F).^54,55^ To determine whether the reduction in global translation we observed upon MGO treatment is also mediated through ISR signaling, we repeated the puromycin incorporation assay in the presence of the ISR inhibitor ISRIB.^56^ We observed that ISRIB partially restored puromycin incorporation relative to MGO treatment alone, indicating a partial rescue of translation (Fig. 4G).

To test whether ISR activation can be directly driven by mRNA glycation, we transfected WT HEK293T cells with our mGFP_MGO_ sensor. Transfection of the glycated RNA resulted in increased eIF2α phosphorylation compared with unglycated mGFP, consistent with mRNA glycation–dependent activation of the ISR, potentially due to impaired ribosome function (Fig. 4H). To directly show that mRNA glycation also triggers the ISR-dependent transcriptional reprogramming, we performed qPCR analyses after transfection with mGFP or mGFP_MGO_. Relative to the unglycated control, mGFP_MGO_ specifically induced the upregulation of the ISR target ATF3, but not the ER stress sensor ATF6, demonstrating that mRNA glycation is sufficient to drive changes in gene expression. Furthermore, treatment with the ISR inhibitor ISRIB during transfection partially rescued the upregulation of this stress gene, confirming that transcriptional reprogramming is mediated through ISR activation (Fig. 4I). Together, these findings indicate that elevated MGO levels induce ribosomal impairment through mRNA glycation, which in turn activates eIF2α and the ISR, leading to transcriptional reprogramming. Thus, MGO acts as a suppressor of translation through two mechanistically distinct but interconnected pathways: (i) direct impairment of highly responsive glycated mRNA templates, and (ii) secondary activation of the ISR via ribosomal impairment, which globally represses cap-dependent translation.

### mRNA glycation activates the ribotoxic stress response (RSR) in a ZAK-dependent manner

While the ISR regulates translation through eIF2α phosphorylation, ribosome impairment can also engage a distinct MAPK signaling cascade known as the ribotoxic stress response (RSR) to activate stress-adaptive responses (Fig. 5A).^9,10,14,15,57^ To investigate whether glycation-induced ribosome impairment also activates the RSR, we exposed WT HEK293T cells to increasing concentrations of MGO and monitored the phosphorylation of p38 and JNK, the hallmarks of RSR activation. Indeed, our results show a dose-dependent upregulation of both p38 and JNK phosphorylation in response to MGO treatment, indicative of RSR-associated MAPK activation (Fig. 5B). This response was also observed in ΔGLO1 cells at lower MGO concentrations (Fig. S4B), confirming that intracellular MGO levels and glycation stress is sufficient to trigger MAPK signaling. Time course analyses revealed the rapid MGO-dependent activation of MAPK signaling, peaking at 1 hour following treatment and persisting for 2 hours (Fig. 5C). This kinetic profile precedes the onset of detectable protein glycation, supporting that these early effects are independent of protein AGEs (Fig. S4C).

**Fig. 5:**
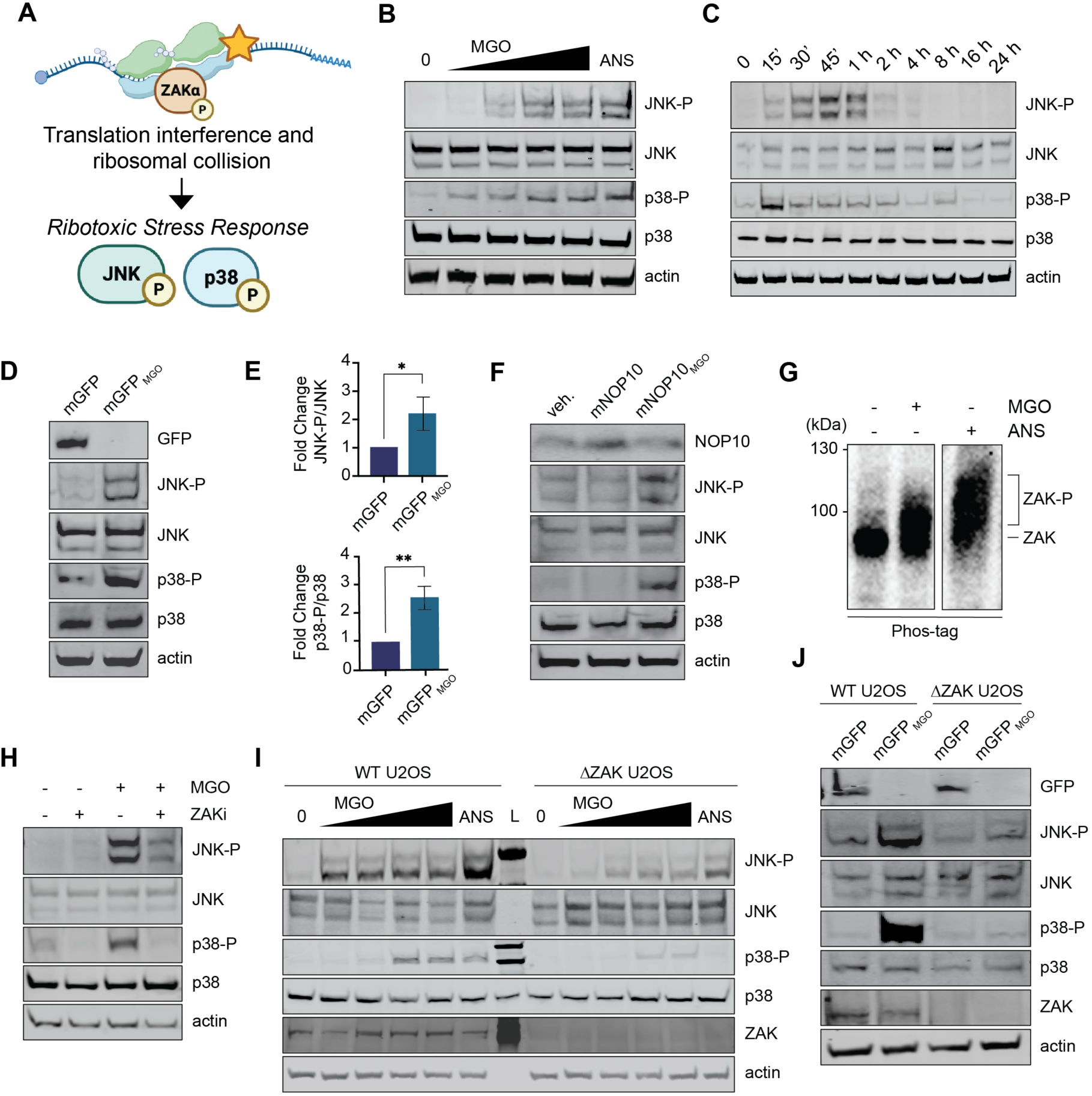
RNA glycation activates the ZAK-dependent ribotoxic stress response. (A) Schematic of ZAK activation upon ribosome collision, leading to the activation of the ribotoxic stress response (RSR). (B) Immunoblot analysis of WT HEK 293T cells treated for 1 hour with 0-0.35 mM MGO or 1 µg/mL anisomycin (positive control for RSR activation) (n=3). (C) Immunoblot analysis of WT HEK 293T cells after treatment with 0.35 mM MGO for the indicated time. () Immunoblot analysis of WT HEK293T cells transfected for 4 hours with either mGFP or mGFP_MGO_ (10 equivalents of MGO per guanosine for 16 hours at 37 °C) (n=3). (E) Quantification of fold change of JNK-P/JNK (n=3) (upper panel) and p38-P/p38 (n=3) (lower panel) from Figure 5D. Statistical significance was assessed by two-tailed unpaired *t* test. (\**P*< 0.0332; \*\**P*< 0.0021). (F) Immunoblot analysis of WT HEK293T cells transfected for 4 hours with a vehicle control, mNOP10, or mNOP10_MGO_ (10 equivalents of MGO per guanosine for 16 hours at 37 °C) (n=3). (G) Immunoblot of Phos-tag gel for ZAK phosphorylation in WT HEK293T cells treated with 0.24mM MGO or 1 µg/mL anisomycin for 1 h (n=3). (H) Immunoblots of WT HEK293T cells pretreated (30 min) with or without ZAK inhibitor (ZAKi, nilotinib, 1 µM), followed by treatment with or without MGO (0.24 mM, 1 h) and ZAK inhibitor (ZAKi, nilotinib, 1 µM) (n=3). (I) Immunoblot analysis of WT and ΔZAK U2OS cells treated for 1 hour with 0-0.24 mM MGO or 1 µg/mL anisomycin (n=3). (J) Immunoblot analysis of WT and ΔZAK U2OS cells transfected for 4 hours transfected with either mGFP or mGFP_MGO_ (10 equivalents of MGO per guanosine for 16 hours at 37 °C) (n=3).

To show the direct impact of RNA glycation on the RSR, we transfected either unglycated mGFP or mGFP_MGO_ and followed p38 and JNK activation by Western blot analysis. Upon transfection of mGFP_MGO_ we observed an increase in p38 and JNK phosphorylation compared to mGFP, reaching levels similar to those observed after MGO treatment, indicating that mRNA glycation can directly activate the RSR (Fig. 5D, E). To demonstrate that RNA-glycation dependent activation of the RSR can also occur following glycation of an endogenous transcript, we focused on NOP10, which was identified by Keth-seq as highly susceptible to glycation (Fig. 2F). We generated NOP10 mRNA (mNOP10) *in vitro* using the same approach as for mGFP and subjected it to *in vitro* glycation to obtain mNOP10_MGO_. Similarly to mGFP_MGO_, mNOP10_MGO_ showed reduced NOP10 translation and induced RSR activation compared to the unglycated mNOP10 upon transfection into cells (Fig. 5F).

To confirm that the observed RSR activation is due to ribosomal interference, we focused on ZAK, the key kinase shown to be activated by autophosphorylation upon ribosome collision and triggers downstream JNK and p38 activation during the RSR.^13^ We first probed for ZAK phosphorylation using Phos-tag gels which slow the migration of phosphorylated proteins.^58^ We observed that MGO induces ZAK activation in WT HEK293T cells (Fig. 5G), detected as a shift in the molecular weight ZAK band corresponding to ZAK phosphorylation, similar to anisomycin treatment (Fig. 5G, S4D). Nilotinib is a commercial compound that was originally developed as an inhibitor for BCR-ABL but is also used as a potent inhibitor of ZAK.^9,59^ Pre-treating WT HEK293T cells with nilotinib (ZAKi), we observed a reduction in MGO-induced p38 and JNK phosphorylation (Fig. 5H) indicating ZAK as a key regulator of MGO-induced RSR. To further validate this finding, we took advantage of a recently developed ZAK knockout U2OS cell line (ΔZAK U2OS)^10^ and compared MGO-induced RSR activation in ΔZAK versus WT U2OS cells. While MGO treatment robustly activated p38 and JNK in WT cells, this activation was substantially reduced in ΔZAK cells (Fig. 5I). Importantly, LCMS analysis revealed comparable levels of RNA glycation in both ΔZAK and WT cells following MGO treatment (Fig. S4E), demonstrating that the impaired stress signaling in ΔZAK cells is not due to reduced RNA glycation but rather reflects the essential role of ZAK in RSR activation. To again demonstrate the direct involvement of RNA glycation in this ZAK mediated pathway, we transfected mGFP_MGO_ into both ΔZAK and WT cells, showing that mGFP_MGO_ was sufficient to induce p38 and JNK phosphorylation in WT but not in ΔZAK (Fig. 5J). Finally, we performed a time course analysis and observed that the activation of p38 and JNK was fully abolished across all time points upon ZAK deletion, further confirming that the RSR activation is ZAK-dependent (Fig. S4G).

Since the RSR is known to mediate downstream cellular outcomes including cell cycle arrest, we next examined whether MGO treatment induces changes in the cell cycle progression in WT HEK293T cells.^9^ As expected, MGO exposure for 1 hour drove the stalling of cells in G0/G1 phase (Fig. S4H), linking glycation-induced RSR activation to cell-cycle checkpoint. Altogether, these data indicate that MGO-induced RNA glycation rapidly triggers the RSR through ribosomal interference and identify ZAK as the key direct sensor of glycation-induced ribosome impairment. Moreover, these results establish RNA glycation as an early and direct trigger of ZAK-dependent RSR signaling, identifying dysregulated glycolysis as a potent inducer of p38 and JNK activation.

### RNA glycation impairs β-cell function via the ribotoxic stress response (RSR)

Next, we investigated the role of RNA glycation–induced RSR activation in diabetes Mellitus (DM), where MGO levels are elevated and linked to disease progression.^22,60^ We focused on JNK, a key mediator of cellular dysfunction in DM.^61–63^ In β-cells, JNK activation suppresses insulin gene expression, promotes inhibitory serine phosphorylation of insulin receptor substrate proteins, attenuates Akt signaling, and lowers glucose-stimulated insulin secretion (GSIS) thereby promoting β-cell failure under metabolic stress.^61,62,64^ Elevated MGO levels were similarly shown to disrupt β-cell function and reduce GSIS, in part via JNK activation.^60,63,65,66^ Because the molecular link between MGO and JNK remains unclear, we investigated whether RNA glycation–induced RSR activation is a possible mechanism connecting metabolic MGO stress to JNK-driven β-cell dysfunction.

Using MIN6 insulinoma cells as a model for pancreatic β-cell function^67^ we first evaluated the levels of RNA glycation. Performing a quantitative LCMS on total RNA extracted from MIN6 cells treated with increasing concentrations of MGO to mimic elevated MGO levels in DM revealed a dose-dependent accumulation of cellular RNA glycation levels, similar to WT HEK293T cells (Fig. 6A). Western blot analysis showed that treatment of MIN6 cells with MGO for 1 hour led to an increased phosphorylation of JNK and eIF2α, confirming MGO-dependent activation of the RSR and ISR in β-cells (Fig. S5A). Next, we examined the role of the RSR and JNK activation in pancreatic β-cell function and co-treated MIN6 cells with MGO and ZAK inhibitor. We observed ZAK-dependent reduction of MGO-induced JNK phosphorylation (Fig. 6B), demonstrating ZAK dependency of the RSR β-cells.

**Fig. 6:**
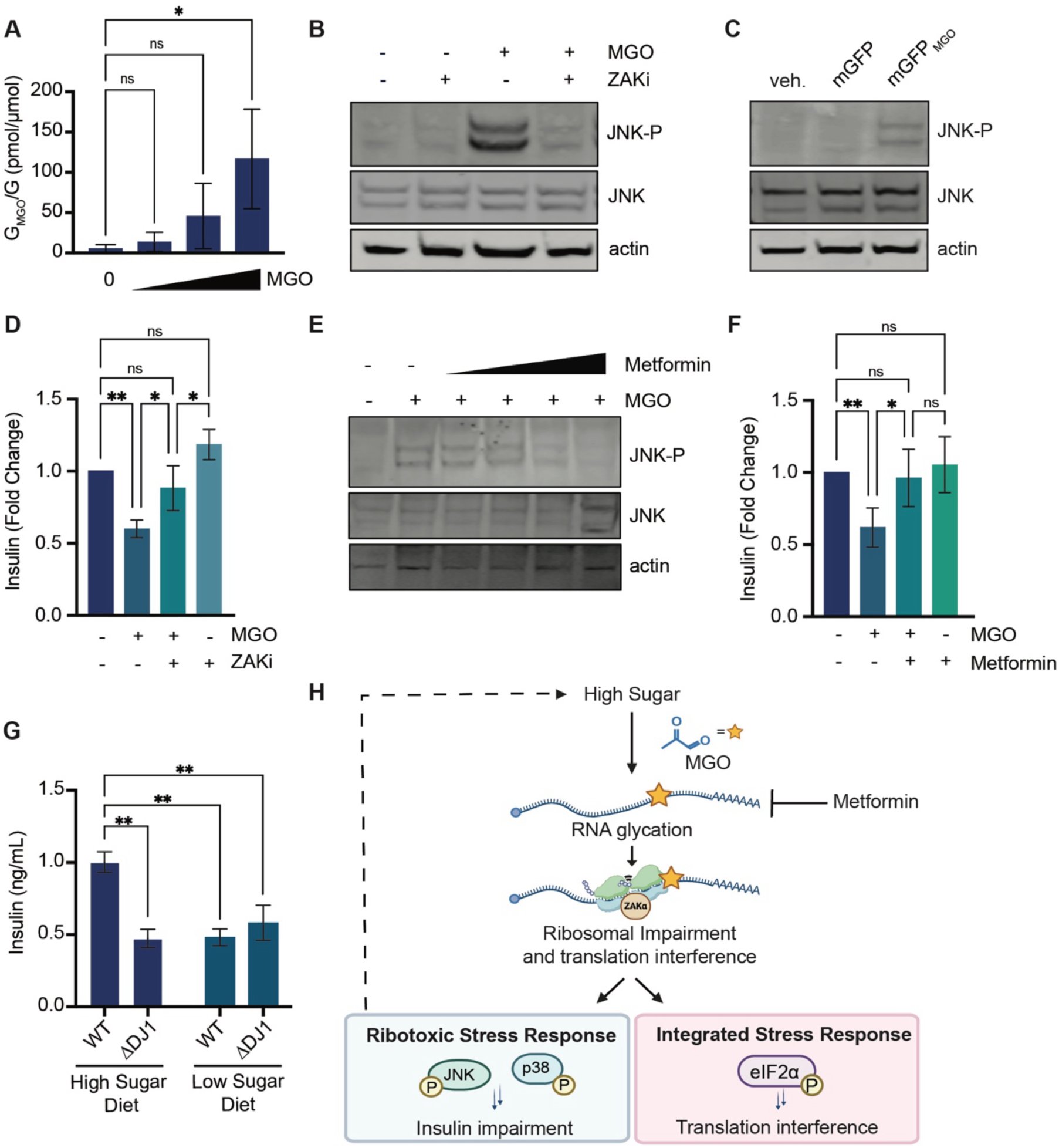
RNA glycation activates the ribotoxic stress response to impair β-cell function and insulin secretion. (A) G_MGO_ levels of total RNA isolated from MIN6 cells treated with 0 – 0.24 mM MGO for 1 hour measured by LCMS (n=3). Statistical significance was assessed using a one-way ANOVA with Dunnett’s multiple-comparison test *(*P<* 0.0332; *\*\*P<* 0.0021; *\*\*\*P<* 0.0002; *\*\*\*\*P<* 0.0001). (B) Immunoblot analysis of MIN6 cells pretreated with DMSO or nilotinib (ZAKi; 1 μM, 30 min), followed by treatment with 0 or 0.24 mM MGO for 1 hour (n = 3). (C) Immunoblot analysis of MIN6 cells transfected with either mGFP or mGFP_MGO_ (10 equivalents of MGO per guanosine for 16 hours at 37 °C) (n=3). (D) Glucose-stimulated insulin secretion (GSIS) in MIN6 cells under high glucose (16.7 mM) stimulation conditions. Cells were preincubated with 0 or 0.24 mM MGO for 1 hour, with or without ZAKi pretreatment (1 μM, 30 min). Data are shown as fold change relative to untreated controls (n=3). Statistical significance was assessed using a Two-way ANOVA with Tukey’s multiple comparison test *(*P<* 0.0332; *\*\*P<* 0.0021; *\*\*\*P<* 0.0002; *\*\*\*\*P<* 0.0001). (E) Immunoblot analysis of JNK phosphorylation in MIN6 cells co-treated with increasing concentrations of metformin (0–50 mM) and with 0.24 mM MGO for 1 hour (n=3). (F) GSIS in MIN6 cells treated with 0.24 mM MGO for 1 hour in the presence or absence of metformin pretreatment (1 mM for 30 min), (n=3), Statistical significance was assessed using a Two-way ANOVA with Tukey’s multiple comparison test *(*P<* 0.0332; *\*\*P<* 0.0021; *\*\*\*P<* 0.0002; *\*\*\*\*P<* 0.0001). (G) Serum insulin levels in WT and ΔDJ-1 mice maintained on low-sugar or high-sugar diets (n=2-3), Statistical significance was assessed using a Two-way ANOVA with Tukey’s multiple comparison test *(*P<* 0.0332; *\*\*P<* 0.0021; *\*\*\*P<* 0.0002; *\*\*\*\*P<* 0.0001). All data are presented as mean ± SD. (H) Model illustrating the pathway in which elevated sugar levels promote RNA glycation, leading to ribosomal impairment, activation of RSR and ISR, translational dysfunction, and reduced insulin secretion.

To assess whether RNA glycation is sufficient to activate the RSR in β-cells, we transfected MIN6 cells with mGFP_MGO_ and showed that mGFP_MGO_ but not mGFP induced JNK phosphorylation, identifying RNA glycation as a direct and proximal trigger of JNK activation in β-cells (Fig. 6C). To determine the role of RNA glycation in β-cell dysfunction, we performed GSIS assays in MIN6 cells. Briefly, cells were treated with MGO for 1 hour and then starved for additional 2 hours. After high-glucose stimulation, the secreted insulin was measured by ELISA. Our results indicate that MGO treatment significantly reduces insulin secretion, consistent with increased JNK activation (Fig. 6D). Notably, pretreatment of cells with the ZAK inhibitor prior to MGO exposure restored insulin secretion to control levels (Fig. 6D), indicating that ZAK-dependent RSR activation mediates β-cell secretory dysfunction under metabolic stress. While MGO-induced JNK activation and reduction of glucose-stimulated insulin secretion have largely been attributed to oxidative stress and AGE–RAGE signaling, our findings reveal a ZAK-centered mechanism in which RNA glycation serves as the initiating event.

Given that elevated MGO is a defining feature of metabolic diseases,^18,19,22,28^ we asked whether pharmacological suppression of MGO could mitigate RNA glycation and its downstream consequences. Metformin is the most widely prescribed first-line therapy for DM and has a long-standing record of improving glycemic control and limiting diabetic complications.^68^ Beyond its AMPK-dependent effects, metformin was suggested to directly target metabolic stress by chemically reacting with MGO to form stable imidazolinone adducts, which can be detected in urine samples from metformin-treated diabetic patients (Fig. S5B).^69^ To assess the effect of metformin on RNA glycation, we incubated total RNA extracted from WT HEK293T cells with MGO in the presence or absence of metformin. RNA was purified, enzymatically digested to mononucleotides, and HPLC analyzed as described above (Fig. 1D). As shown in Fig. 1D, MGO treatment induced a selective reduction in the guanosine peak and generation of new peaks corresponding to glycation adducts. Importantly, co-treatment with MGO and metformin markedly reduced RNA glycation by restoring the guanosine peak and reducing the modified guanosine peaks (Fig. S5C). To examine whether metformin modulates glycation-induced RSR activation *in cellulo*, we treated MIN6 cells with increasing concentrations of metformin and MGO for 1 hour. Western blot analysis showed that while MGO induces JNK activation, co-treatment with metformin and MGO dose-dependently reduced JNK phosphorylation, consistent with reduced RSR activation in β-cells (Fig. 6E).

To assess whether metformin can counteract RSR-mediated β-cell dysfunction, we performed GSIS assays in MIN6 cells treated with metformin and MGO for 1 hour. We measured insulin secretion levels by ELISA and observed that MGO treatment reduces insulin secretion, whereas co-treatment with metformin restored insulin release, indicating that metformin protects β-cells from glycation-induced secretory defects (Fig. 6F). Together, these findings demonstrate that metformin attenuates MGO-induced JNK activation and alleviates RNA glycation–associated defects in insulin secretion, revealing a previously unappreciated mechanism by which metformin may limit DM progression and related complications. Finally, we examined whether DJ-1–mediated attenuation of RNA glycation (Fig. 1D-E) also regulates insulin secretion *in vivo* by measuring serum insulin levels of WT vs DJ-1 knockout mice under different dietary inputs. Briefly, WT and DJ-1 knockout mice were subjected to low- or high-sugar diets for 12 weeks, and serum samples were subsequently collected for insulin quantification. Indeed, DJ-1 knockout mice fed with a high-sugar diet exhibited significantly reduced circulating insulin levels compared with WT mice controls (Fig. 6G). This result is consistent with our observation that DJ-1 loss leads to increased MGO levels, which regulate JNK activation via ribosomal dysfunction and impair β-cell function. In contrast, mice maintained on a low-sugar diet showed no difference in insulin levels between genotypes, likely reflecting minimal insulin demand under non-stimulatory conditions. These results suggest that DJ-1 protects β-cells from MGO stress *in vivo* by limiting MGO accumulation and preventing RNA glycation, downstream RSR activation and JNK-mediated secretory dysfunction.

Altogether, these findings reveal a previously unrecognized mechanism by which metabolic stress impairs translation and β-cell function through RNA glycation. Under conditions of high glycolytic flux, elevated MGO levels modify highly susceptible mRNAs. Glycation adducts on these mRNAs act as a sensor and induce ribosomal impairment which activates two distinct signaling pathways: First, the ISR which leads to cap-dependent downregulation of translation and translational reprogramming. Secondly, the ZAK-dependent RSR, leading to p38 and JNK phosphorylation. In β-cells, ZAK-dependent JNK activation drives a decreased insulin secretion (Fig. 6H). Importantly, ZAK inhibition restores insulin secretion, highlighting a potential therapeutic target for metabolic dysregulation. Moreover, the capacity of metformin to reduce MGO accumulation, prevent RNA glycation and JNK activation and rescue β-cell function underscores a novel mechanism how metformin can preserve β-cell function.

## Discussion

Excessive dietary sugar consumption and type 2 diabetes mellitus (DM) are characterized by elevated levels of reactive carbonyl species such as methylglyoxal (MGO), driving the formation of advanced glycation end products (AGEs) on biomolecules.^28,60^ Thus far, protein and DNA glycation have been established as contributors to diabetic pathology through oxidative stress and AGE–RAGE signaling. While free glycated guanosine (CEG) adducts have been shown to accumulate extracellularly in patient with DM^33,34^ and preliminary reports showed that RNA glycation can interfere with translation,^33,35,46^ their regulation, the mechanism governing the translation interference as well as the downstream biological impact of RNA glycation in cells is unknown. We propose a model in which elevated glycolytic flux increases intracellular MGO, leading to rapid glycation of highly susceptible mRNAs which is dynamically regulated by DJ-1 and the glyoxalase system. Glycation of these mRNAs leads to ribosomal impairment and decreased translation and act as a sensor to activate two coordinated stress pathways: (i) the ISR, which suppresses global cap-dependent translation and promotes adaptive reprogramming, and (ii) the ZAK-dependent RSR, which drives β-cell dysfunction through JNK phosphorylation. Notably, JNK activation is attenuated by metformin, offering a new mechanistic role of how metformin protects β-cells function.

Our characterization of RNA MGO-glycation expand the canonical epitranscriptome and highlighting it as a novel, highly regulated non-enzymatic RNA modification (NECM) that accumulates under metabolic stress.^5^ Unlike enzymatic RNA modifications, which are installed by dedicated “writer” enzymes that recognize often specific sequence motifs,^70^ NECMs occur spontaneously. However, our results show that RNA glycation is not distributed stochastically, but displays a degree of selectivity, that appears to be driven not by defined RNA sequence motifs, but by the structure, half-life and abundance of the mRNA.^42^ Although RNA glycation arises non-enzymatically, our results show that its removal is dynamically regulated by DJ-1, reminiscent of how RNA “erasers” such as FTO and ALKBH5 regulate canonical marks such as m⁶A.^3^

More broadly, NECMs on RNA influence metabolic processes and cellular function much like their enzymatic counterparts,^5^ yet only a limited subset has been characterized.^9,10,14,57,71^ These include modifications induced by UV light, ROS and alkylation, which were shown to impair translation and lead to ribosome stalling and collisions,^5,9,10,13–15,57,71,72^ effects that have also been recently reported for canonical modifications such as m⁶A.^52^ Pioneering studies using UV light, such as during sunburn in keratinocytes, and ROS generated under conditions like high-fat, high-sugar diets have demonstrated how these lead to ribosome impairment, which is translated into activation of the ISR and RSR, thereby regulating translation, metabolic adaptation, and cell fate decisions.^9,10,13–15,57,71^ Extending this framework, we identify RNA glycation as an additional and physiologically relevant driver of ribosome dysfunction under metabolic stress, activating both the ISR and ZAK-dependent RSR. While additional drivers of MGO-induced stress responses such as MGO-induced ROS, glycation of other biomolecules or ER stress cannot be fully excluded,^63,73^ our data recapitulate all downstream signaling events by the direct transfection of glycated mRNA (mGFP_MGO_), demonstrating that RNA glycation is sufficient for activating the RSR and ISR. We also observe RNA glycation-specific effects at short time scales (30 minutes to 1 hour), substantially prior to the appearance of protein glycation, which occurs after 2-4 hours. These findings are consistent with the prevailing hypothesis in the literature that non-enzymatic RNA modifications function as early sensors of cellular stress and regulator of cell fate decisions.^5,48^

Given the pronounced metabolic perturbations characteristic of DM, we therefore propose that intracellular RNA glycation is not merely a biomarker of the disease^34^ but actively contributes to diabetic progression and complication by engaging both the ISR and the RSR via ribosomal impairment. Activation of JNK contributes to the development of insulin resistance and β-cell dysfunction, hallmarks of DM, and is associated with metabolic inflammation that exacerbates disease progression and diabetic complications.^61^ Interestingly, the ISR is classically viewed as a protective stress response that promotes adaptive translational reprogramming,^53,61,74^ but its activation has also been associated with DM comorbidities.^75,76^ This offers a novel mechanism whereby RNA glycation signals pathways that promote DM and its complications, unifying several previous observations.^62–64^

Building on this concept, we further examined the impact of metformin on RNA glycation, a first-line antihyperglycemic therapy with reported anti-glycation activity.^69^ Consistent with clinical and experimental studies showing that metformin reduces circulating MGO levels and AGE burden,^68,77,78^ co-treatment of metformin with MGO markedly decreases RNA glycation *in vitro* and attenuates RSR signaling. Importantly, metformin restored insulin secretion under MGO-induced stress, functionally linking suppression of RNA glycation and RSR activation to β-cell recovery. While metformin exerts additional effects on β-cell function, including AMPK-dependent pathways,^79^ our data suggest that mitigation of RNA glycation is sufficient to dampen ribosomal stress signaling and rescue insulin secretion. These findings position metformin not only as a metabolic regulator but also as an indirect modulator of RNA integrity and ribosome-mediated stress responses in β-cells.

Cells rely on endogenous detoxification systems to alleviate glycation stress, primarily the glyoxalase pathway (GLO1/2) and DJ-1, both known to support β-cell survival and insulin secretion.^37,80^ Here, we show that both GLO1 and DJ-1 regulate RNA glycation levels using knockout cell lines of GLO1 and DJ-1 as well as DJ-1–deficient mice. While our *in vitro* data indicates that DJ-1 can directly lower RNA glycation levels, it is also possible that RNA glycation reflects a rapidly reversible process in which glycated and unglycated states exist in equilibrium, and that MGO quenching by DJ-1 shifts this balance toward the unglycated state.^25,39,81^ Consistent with our model, DJ-1–deficient mice exhibited reduced insulin secretion, providing *in vivo* evidence that DJ-1–mediated control of RNA glycation is a critical upstream regulator of the ZAK–JNK signaling axis to regulate insulin secretion.

In conclusion, our results expand the scope of non-enzymatic glycation from proteins and DNA to RNA, revealing that RNA itself is an important target of metabolically driven chemical modifications that are dynamically regulated by cellular detoxification pathways. This positions RNA glycation within the broader landscape of the epitranscriptome, highlighting that non-enzymatic RNA modifications can impact RNA function and cellular responses to stress. These findings open new avenues for understanding how metabolic stress reshapes RNA function and suggest that targeting RNA glycation and its downstream signaling may provide therapeutic opportunities in metabolic disease.

### Limitation of the study

While our work defines a mechanistic link between RNA glycation, translational stress, and β-cell dysfunction, the mapping strategy employed here relies on kethoxal as a chemical mimic for glycation and may not fully capture the spectrum, kinetics, or sequence specificity of endogenous MGO-mediated RNA modifications under physiological metabolic stress. Transcriptome-wide identification of native RNA glycation sites covering all spectrum of glycation adducts in cells experiencing elevated glycolytic flux would be necessary to determine the sequence or structural determinants of RNA susceptibility to glycation; however, unbiased approaches to map endogenous RNA glycation in living cells are currently unavailable. While we have demonstrated RNA glycation-dependent ribosome collision *in vitro*, *in vivo* polysome profiling was not performed in this study as MGO glycation was shown to rearrange and form protein–nucleic acid cross-linking,^7^ which interferes with the RNase digestion steps required for accurate polysome fractionation. Consequently, standard polysome profiling approaches are not likely to be compatible with MGO-treated samples.

## Experimental Models

### Cell lines

MIN6 cells were a kind gift from Danwei Huangfu Lab (MSKCC). Wild-type U2OS and the ΔZAK U2OS cells were a kind gift from Simon Bekker-Jensen Lab (University of Copenhagen). The HEK293T cells with GLO1 (ΔGLO1) or DJ-1 (ΔDJ-1) deletion were received from the Jim Galligan Lab (University of Arizona). HEK293T and U2OS cell lines were cultured in high glucose DMEM supplemented with 10% FBS, 2 mM glutamine, 10 U/mL penicillin, and 10 μg/mL streptomycin. MIN6 cells were cultured in high glucose DMEM supplemented with 15% FBS, 2 mM glutamine, 10 U/mL penicillin, 10 μg/mL streptomycin, and 50 μM β-mercaptoethanol.

### Animal models

All animal procedures were approved by the Columbia University Institutional Animal Care and Use Committee (IACUC) (protocol #AABV8661). Mice were housed in the Columbia University Russ Berrie Medical Science Pavilion under controlled conditions (12 h light/dark cycle; 50–60% humidity). Eight- to ten-week-old male and female C57BL/6J (JAX #000664) and DJ-1⁻/⁻ (JAX #006577) mice were obtained from The Jackson Laboratory. Animals had *ad libitum* access to water and standard chow (PicoLab Rodent Diet 20, #5053) during breeding and non-experimental maintenance.

For experimental studies, mice assigned to the low-sugar diet were fed *ad libitum* with diet #D12450K (Research Diets, Inc.) and regular water, whereas mice assigned to the high-sugar diet received diet #D12492 (Research Diets, Inc.) supplemented with *ad libitum* high-fructose/high-glucose drinking water (23.1 g/L D-fructose and 18.9 g/L D-glucose). Animals were maintained on these diets for 12 weeks prior to euthanasia by CO₂ asphyxiation. For plasma collection, blood was obtained by cardiac puncture and transferred into tubes containing 10 µL of 0.5 M EDTA. Samples were incubated on ice for 15 min and centrifuged at 500×g for 10 min at 4 °C. The plasma supernatant was collected and stored at −80 °C until analysis. Livers were harvested immediately after euthanasia, rinsed in cold PBS, and either snap-frozen in liquid nitrogen.

## Method Details

### LCMS ANALYSIS OF RNA GLYCATION

#### Synthesis of LCMS standards

Isotopically labeled standards of N2-(1-carboxyethyl)guanosine (CEG) and its cyclic form (cyclic CEG, also referred to as cMGO-guanosine or c-MGO-G) were synthesized similar to previously described methods (Fig. S1A). ^32^ In brief, 10 μM of Guanosine-^15^N_5_ 5′-monophosphate was incubated with Shrimp Alkaline Phosphatase for 3 hour at 37 °C until dephosphorylation was complete. The identity of ^15^N-Guanosine (^15^N-G) was confirmed by LCMS (Fig. S1B). ^15^N-G was incubated with 5 eq MGO for 4 hours and reaction mixture was purified via flash high-pressure reverse-phase liquid chromatography on an Agilent 1200 series instrument with Agilent Zorbax SB-18C column (5 μm, 9.4 mm × 150 mm, gradient 1-5%B in 5 minutes, 5%B for 45 minutes, A: water + 0.01% TFA B: 90% MeCN + 0.01% TFA). For LCMS/MS analysis, the reaction mixture was diluted 1:100 or 1:10 with 0.1% aqueous formic acid. The identities of ^15^N-CEG and ^15^N-c-MGO-G (collectively named ^15^N-G_MGO_) were confirmed by comparing MS/MS fragmentation patterns to expected fragment masses based on the chemical structures (Fig. S1C).

#### MS fragmentation analysis

MS and MS/MS analyses were performed on a QTRAP 6500 mass spectrometer (SCIEX) equipped with a Turbo Spray IonDrive source. For direct infusion experiments, working solutions were prepared in 80% methanol in nanopure water and infused directly into the mass spectrometer. Negative ion mode was used for all analyses. Precursor ions of m/z 282 (guanosine, [M-H]−) and m/z 354 (CEG and c-MGO-G, [M-H]−) were selected for MS/MS fragmentation. For ^15^N-labeled standards, m/z 287 (^15^N-G) and m/z 359 (^15^N-G_MGO_) were monitored. Collision energy (CE) and declustering potential (DP) were optimized for each transition.

#### LCMS/MS method development

Chromatographic separation was achieved using an Atlantis dC18 column (100 Å, 3 μm, 2.1 mm × 50 mm, Waters) maintained at ambient temperature. The mobile phase consisted of 0.1% formic acid in water (Buffer A) and 0.1% formic acid in acetonitrile (Buffer B) at a flow rate of 0.4 mL/min. The gradient program was as follows: 0–3.0 minutes, 3-10% B; 3.0–4.0 minutes, 10-100% B; 4.0–5.5 minutes,100% B; 5.5–6.0 minutes, 100% B returning to 3% B. The injection volume was 10 μL for guanosine and 12 μL for CEG and c-MGO-G. Detection was performed in negative ion multiple reaction monitoring (MRM) mode. The following transitions were monitored: guanosine 282→150 (quantifier) and 282→133; CEG and c-MGO-G 354→178 (quantifier) and 354→204; ^15^N-guanosine 287→155 (quantifier) and 287→137; ^15^N-CEG 359→209 and 359→183 (quantifier). Consistent with previously reported DNA (deoxyguanosine)-AGE products^32^, CEG and c-MGO-G are also at reversible equilibrium. Peak identity assignments were based on polarity considerations; the more polar CEG (due to the carboxylic acid group) eluted earlier (at approximately 2.1–2.2 minutes), while the less polar (c-MGO-G) eluted later (at approximately 3.1–3.4 minutes, Fig. S1F). These elution patterns are consistent with previous reports of DNA-AGE products.^32^

#### Titration and linear dynamic range determination

Titration analyses were performed to establish the linear dynamic range and determine optimal internal standard concentrations for biological sample analysis. For ^15^N-guanosine, linear regression analysis demonstrated linearity between injected amount (0–600 pmol) and area under curve (AUC) with R2 = 0.99 for the 287→155 transition (Fig. S1D). For ^15^N-G_MGO_, linearity was established for the 359→183 transition at both 2.38 minutes (CEG) and 3.38 minutes (c-MGO-G) retention times (R2 = 0.96, Fig. S1E). Based on these analyses, 600 pmol of ^15^N-G and 6 pmol of ^15^N-G_MGO_ were determined to be optimal for spiking into biological samples, maintaining analyte-to-standard area ratios within 1:10.

#### Biological sample preparation for G_MGO_ quantification

Cells were treated with MGO as indicated and total RNA was extracted using TRIzol. For the comparison of mRNA to total RNA, RNA was enriched for polyA RNA by using Dynabeads™ Oligo(dT)_25_. RNA was fractionated using Cytoplasmic and Nuclear RNA purification kit. All RNA was subjected to in-column DNAse digestion and cleaned up using RNA Clean & Concentrator-5. After digestion of the RNA using the Nucleoside Digestion Mix for 16 hours at 37 °C, the samples were lyophilized. Mice liver tissues of WT and ΔDJ-1 mice, fed with a low sugar diet, were homogenized using the TissueLyser II (Quiagen) and the RNA was extracted using TRIzol and further processed as described above.

Given the low abundance of CEG and c-MGO-G observed in preliminary analyses, samples were reconstituted in 50 μL of a mixture containing 97% of 0.1% formic acid in water and 3% of 0.1% formic acid in acetonitrile to maximize detection sensitivity. ^15^N-G_MGO_ internal standard (10 μL of 3 μM solution) was added to each sample. Samples were briefly sonicated using a Branson Ultrasonics CPXH Series Ultrasonic Cleaning Bath prior to analysis.

#### Biological sample preparation for guanosine quantification

For guanosine normalization, 10 μL of the reconstituted biological sample was diluted with 990 μL of the same solvent mixture (97% of 0.1% formic acid in water and 3% of 0.1% formic acid in acetonitrile) and spiked with 10 μL of 2.5 mM ^15^N-guanosine. This 1:100 dilution was necessary to bring guanosine concentrations within the established linear dynamic range.

#### Quantification of CEG and c-MGO-G

Peak integration and quantification were performed using Analyst software (SCIEX). Area ratios were calculated as AUC of analyte (CEG: 354→178 at approximately 2.1 minutes; c-MGO-G: 354→178 at approximately 3.2 minutes) divided by AUC of ^15^N-CEG internal standard (359→183). The molar amount (pmol) of CEG and c-MGO-G was calculated by multiplying the area ratio by the amount of internal standard (30 pmol total in the 60 μL reconstituted sample volume). Molar amounts were normalized to total guanosine content to account for variations in RNA input.

#### Quantification of guanosine

Guanosine quantification was performed using the 282→150 transition with ^15^N-guanosine (287→155) as internal standard. Area ratios were multiplied by 25 nmol (the amount of ^15^N-guanosine in the analyzed volume) to calculate the molar amount of guanosine. Since 10 μL of the 60 μL total reconstituted sample was used for analysis, the calculated amount was multiplied by 6 to determine total guanosine content per sample.

### EXPRESSION AND PURIFICATION OF DJ-1

The expression and purification of the wild-type were carried out as previously described.^29^ Briefly, pET3a-His-DJ-1 plasmid was transformed into E. coli (BL21) cells and grown for 5 hours at 37 °C before induction with 500 μM IPTG for 3 hours at 37 °C. The bacterial pellet was lysed (50 mM Tris pH 7.5, 100 mM NaCl, 1 mM DTT, 10% glycerol) before being pelleted at 20,000 g for 20 minutes. The His6-tagged DJ-1 was isolated using Ni^2+^ beads and further purified via size exclusion chromatography. The protein purity was verified by SDS-PAGE and LCMS analyses. The protein concentration was determined using 280 nm wavelength on a NanoDrop 2000c (Thermo Scientific).

### *IN VITRO* GLYCATION ASSAYS

Total RNA was extracted from WT HEK293T cells using TRIzol reagent (Thermo Fisher Scientific) following the manufacturer’s protocol. For MGO coincubation experiments, 5 μg of total RNA was incubated in a total volume of 50 μL in three conditions for 16 hours at 37 °C: (i) nuclease-free water (ii) 1.2 mM MGO in nuclease-free water (iii) 1.2 mM MGO and 4 μM recombinant DJ-1 purified in house (described above). For RNA pre-glycation experiments, 5 μg of total RNA was incubated in 50 μL of 1.2 mM MGO in nuclease-free water for 16 hours at 37 °C before free MGO was quenched by supplementation with 25 mM Tris. 4 μM recombinant DJ-1 was added to the pre-glycated RNA and incubated for 16 hours at 37 °C. For metformin coincubation experiments, 5 μg of total RNA was incubated in a total volume of 50 μL in three conditions for 16 hours at 37 °C: (i) nuclease-free water (ii) 1.2 mM MGO in nuclease-free water (iii) 1.2 mM MGO and 1.2 mM metformin in nuclease-free water. RNA samples were digested with 1 μL of nuclease in the supplied digestion buffer (NEB Nucleoside Digestion Mix, M0649) for 16 hours at 37 °C. Digested samples were analyzed using analytical reversed-phase HPLC (RP-HPLC) on Agilent 1260 Quadrupole LC/MS equipped with an Agilent Zorbax C18 column (3.5 µm, 4.6 x 150 mm), employing 0.1% formic acid (FA) in H_2_O (solvent A), and 90% acetonitrile, 0.1% FA in H2O (solvent B), as the mobile phases at a flow rate of 0.5 mL/min, gradient 0-2% B in 2 minutes, 2-7% B in 5 minutes, 7-10% B in 22 minutes.

### REANALYSIS OF PREVIOUSLY PUBLISHED KETH SEQ ANALYSIS

The per-base reactivity score file for each transcript (“GSE122096_HeLa_kethoxal_vs_no-treat.shape.out.txt.gz”) was downloaded from the corresponding GEO repository, comprising a total of 3,522 transcripts. The genome assembly used in the original study was Ensembl hg38 (version 80), which was further employed for all downstream analyses. Unless otherwise specified, all analyses were conducted in R using custom scripts and functions.

Transcripts with nonzero reactivity scores at one or more nucleotide positions were selected for further analysis, resulting in a total of 3,489 transcripts.

A custom R function was developed to integrate the reactivity score files with the corresponding cDNA FASTA file (derived from the same genome assembly) to:

- Verify that transcript lengths matched between the reactivity and cDNA files,
- Count the total number of guanines (G) per transcript, and
- Quantify the number of highly reactive guanines, defined as those with a reactivity score greater than 0.5.

After filtering for transcripts with consistent length between the reactivity and cDNA files, 3,339 transcripts remained. The summary output was used to plot the distribution of transcripts across bins representing the percentage of glycated Gs, calculated as the ratio of highly reactive Gs to total Gs per transcript. These transcripts were further analyzed using the ggscatter from the ggpubr R package to assess the Pearson correlation between transcript length and the number of Gs with a reactivity score > 0.5.

Next, the genomic context annotations of highly reactive bases were characterized using the GenomicFeatures package. A TxDb object was generated from the Homo_sapiens.GRCh38.80.gtf annotation file. Non-overlapping transcript regions were then annotated using the cdsBy, fiveUTRsByTranscript, and threeUTRsByTranscript functions. Highly reactive G were then mapped to one of these categories. The counts were converted to percentages and visualized as a pie chart. Notably, highly reactive Gs located within transcripts lacking defined CDS regions were classified as “other.”

Finally, the distribution of all reactivity scores for each nucleotide (G, A, T, and C) was examined globally, across all transcripts. This analysis confirmed that high reactivity scores were skewed specifically toward guanines. To compare transcript stability, half-life data from publicly available Actinomycin D assay in HeLa cells was utilized (GEO:GSM8410737). A Kolmogorov–Smirnov test was performed to evaluate differences in the distribution of log2-transformed half-lives between transcripts containing at least one highly reactive base (reactivity > 0.5) and those without reactive sites.

For Translation efficiency analyses, Ribo-seq and bulk RNA-seq datasets were obtained from GSM2817679. TPM-normalized (Transcript Per Milion) expression values for the untreated samples in the study were downloaded for two replicates. For each dataset, TPM values were averaged across duplicates.

Transcripts were retained for downstream analysis if they exhibited TPM > 1 in either the Ribo-seq or bulk RNA-seq dataset. Translational efficiency (TE) was calculated for each transcript as the ratio of Ribo-seq TPM to bulk RNA-seq TPM.

Log2-transformed TPM values (log2(TPM+1)) or ratio of those for TE, were compared between transcripts classified as kethoxal-reactive and non-reactive. Statistical significance was assessed using the two sided Wilcoxon rank-sum test implemented in R.

### HPG ASSAY

Cells were cultured in RPMI (without methionine), 1% PSG, and 10% FBS for 1 hour at 37 °C in a flat-bottom 96-well plate. 100 μM HPG was added per well and incubated for 1 hour at 37 °C. After washing with PBS, 100 μL of trypsin was added per well and incubated for 5 minutes before adding 100 μL of media. The cells were transferred to U-bottom 96-well plate and washed with PBS before fixation with 200 µL/well of 3.7% Paraformaldehyde, for 15 minutes at room temperature. The cells were washed with 100 µL/well of 3% BSA (or 3% FBS) in PBS. 100 µL/well of 0.5% Triton® X-100 in PBS were added and incubated for 20 minutes at room temperature before adding the 1X Click-iT® HPG reaction cocktail. The reaction cocktail was composed by 1× Click-iT HPG reaction buffer, copper sulfate, Alexa Fluor 488, and 10× Click-iT HPG buffer additive at a ratio of 2000:43:0.5:10, respectively. The cells were washed with 3% BSA in PBS and 100 µL/well Click-iT® reaction cocktail was added before incubating for 30 minutes at room temperature in the dark. The reaction cocktail was removed and the cells were washed with 100 µL/well Click-iT® reaction rinse buffer before flow cytometry analysis.

### WESTERN BLOTTING

Cells were lysed in RIPA buffer (1x PBS, pH 7.5, 0.5%w/v sodium deoxycholate, 1% NP-40, 0.1% SDS, protease and phosphatase inhibitors). Protein levels were normalized and separated on a 10% Bis-Tris gel with MOPS running buffer and transferred to a PVDF membrane using a Criterion blotter system (Bio-Rad). Membranes were blocked with 5% BSA and proteins of interest were then probed using the appropriate antibody. Membranes were then visualized on a Li-Cor Odyssey CLx fluorescent imaging system. Background, brightness and contrast of the Western Blot images were adjusted using Odyssey CLx fluorescent imaging system, ImageJ and Adobe Illustrator.

### PHOS-TAG SDS-PAGE

For Phos-tag gel immunoblotting, cells were quickly rinsed once with warm PBS and lysed directly in the well using ice-cold lysis buffer (25 mM Tris-HCl, pH 7.6, 150 mM NaCl, 1% sodium deoxycholate, 1% NP-40, 10 mM sodium pyrophosphate, 10 mM β-glycerophosphate, 1 mM TCEP, 170 nL/mL benzonase, and 1× EDTA-free protease and phosphatase inhibitors). Lysates were kept on ice for 10 min and clarified by centrifugation at 8,000 × g for 5 min at 4 °C. After centrifugation, supernatants were collected and boiled in Laemmli SDS sample buffer containing 5% DTT at 95 °C for 10 min. Samples were then resolved by 8% SDS–PAGE containing 10.7 μM Phos-tag acrylamide (Wako, AAL-107) and 21.3 μM MnCl₂. EDTA-free pre-stained protein markers (Apex Bio, F4005) was used. Electrophoresis was performed in 1× Tris–glycine–SDS running buffer. Gels were then washed twice for 10 min each in transfer supplemented with 1 mM EDTA, followed by two additional 10-min washes in transfer buffer without EDTA, and proteins were transferred to 0.2 μm PVDF membranes overnight at 35 V at room temperature. Membranes were blocked with 5% nonfat milk in TBS-T for 1 hr at room temperature with gentle rocking, washed in TBS-T, and incubated with ZAK antibody (Bethyl Laboratories, A301-993A) overnight at 4 °C with gentle rocking. Blots were washed with TBS-T three times for 10 min each, incubated with secondary antibodies diluted in blocking buffer for 1 hr at room temperature with gentle rocking, washed again with TBS-T six times for 10 min each, developed using SuperSignal™ West Atto Ultimate Sensitivity Substrate (Thermo Fisher Scientific, A38556), and imaged on a Bio-Rad ChemiDoc imaging system.

### PUROMYCIN INCORPORATION ASSAYS

Cells were treated as shown in the figures followed by the addition of puromycin (10 µg/ml) to the medium for 5-10 minutes. Cells were washed with ice-cold PBS and immediately lysed. Western blotting with anti-puromycin antibodies was performed to visualize puromycin incorporation into nascent polypeptide chains.

### mRNA *IN VITRO* TRANSCRIPTION AND TRANSFECTION

Plasmid for GFP mRNA production was obtained from addgene. pGEM4Z-T7-5’UTR-EGFP-3’UTR-A64 was a gift from Christopher Grigsby & Molly Stevens (Addgene plasmid # 203348). The plasmid was linearized with SpeI at 37 °C for 30 minutes and purified by ethanol precipitation. For NOP10 mRNA production, a geneblock was amplified using appropriate primers (Table S3). *In vitro* transcription of the linearized template or amplified PCR DNA template was conducted using a T7 RNA polymerase–based high-yield transcription system (AmpliScribe T7 Flash/High Yield). Following a 2 hour incubation at 42 °C, template DNA was degraded by RNase-free DNase, and RNA was recovered by ethanol precipitation and resuspended in nuclease-free water. Capping of purified RNA was performed using the Vaccinia Capping System. The enzymatic addition of 2′-N3-2′-dATP (1 mM) to the poly(A) tail of GFP mRNA (∼3–4 μg) was performed in the presence of yeast poly(A) polymerase (600 U) in 1 × PAP reaction buffer for 20 minutes at 37 °C followed by ethanol precipitation overnight and washing with 70% ethanol. The click reaction was performed with azido-modified mRNA (500 ng) and DBCO-TAMRA (50 μM) for 45 minutes at 37 °C followed by ethanol precipitation overnight and washing with 70% ethanol. Yield and quality were assessed using a NanoDrop™ One/OneC Microvolume UV Vis Spectrophotometer (Thermo Scientific) and gel electrophoresis respectively. For transfections, 2.5 µL of Lipofectamine MessengerMAX and 5 µg mRNA were incubated in separate 125 µL Opti-MEM reduced serum medium for 10 minutes, combined and incubated for 5 minutes, then added dropwise to a 70% confluent 6 cm plate of cells.

### LIVE CELL IMAGING

WT HEK293T cells in DMEM+10% FBS media were seeded on 66.66 µg/mL collagen I-coated 1cm^2^ polymer coverslip bottom wells (Ibidi, Germany) at 150,000 cells/well overnight. The cells were transfected with mGFP-TAMRA and mGFP-TAMRA_MGO_ (0.5 µg per well) using Lipofectamine MessengerMax Reagent. Two hours post transfection cells were washed once with PBS, replenished with fresh DMEM, and were imaged every 15 minutes for 8 hours using an Orca-Fusion BT CMOS camera and Zeiss AxioObserver.Z1 microscope equipped with a 20x/0.8NA objective lens. The FITC filter was used for green fluorescence detection, TRITC filter for the red, and transmitted light for the brightfield. A 9-slice z-stack for a total height of 14 µm was obtained to confirm RNA localization. Cells were maintained at 37 °C and 5% CO_2_. Images were analyzed using FIJI. A top hat filter was used on the transmitted light image to segment the cells. Then the intensity of the FITC channel was measured at each time point to get the average fluorescence per cell at every timepoint.

### *IN VITRO* COLLISION ASSAY

*In vitro* translation reaction was performed as previously described with some modification.^52^ To setup an *in vitro* system to visualize monosome and collided ribosome footprints, GFP mRNA was prepared using α-[32P]-CTP and 3′-O-Me-m7G(5’)ppp(5’)G RNA Cap Structure Analog along with cold ATP, UTP, CTP, and GTP using HiScribe T7 High Yield RNA Synthesis Kit. The synthesized mRNA was purified using 1x volume of RNAClean XP beads according to the manufacturer’s instruction and quantified using Qubit Broad RNA Assay Kit. To test if mRNA glycation can induce ribosome collision *in vitro*, 640 ng of α-[32P]-CTP labeled GFP mRNA was treated with the indicated concentration of MGO for 6 hours at 37 °C. The glycation reaction was quenched with 2 mM arginine. Glycated mRNA was purified using RNAClean XP beads and quantified using Qubit Broad RNA Assay Kit. 2 pmol of the purified mRNA was used for 12.5 μL *in vitro* translation reaction in HeLa lysate (ThermoFisher) for 25 minutes at 30 °C. At the end of the experiment, we stopped the reaction by adding 200 μM emetine. The reaction lysate was subjected to partial digestion using Micrococcal nuclease (1400 activity unit) in the presence of 15mM CaCl_2_ for 45 minutes at room temperature to obtain ribosome footprints. Ribosome footprints were purified using TRIzol according to the manufacturer’s instruction and ran on 15% TBE-urea gel followed by transferring to a nylon membrane (Amersham Hybond-N+, RPN203B) for autoradiography. Radioactive signals were visualized using an Amersham Typhoon 5. To confirm that the appearance of ribosome footprints is translation dependent, each reaction was carried out with or without the addition of 0.5 μM recombinant eIF4E.

### RNA-SEQ and ISR ANALYSIS

ΔGLO1 HEK293T Cells were treated with 60 µM methylglyoxal (MGO) for 4 h, with untreated cells serving as controls. Following treatment, cells were directly lysed in TRIzol reagent (Invitrogen) and total RNA was extracted according to the manufacturer’s protocol. Briefly, cells were lysed directly in TRIzol, followed by chloroform phase separation and isopropanol precipitation. RNA was washed with 75% ethanol and resuspended in RNase-free water. After RiboGreen quantification and quality control by Agilent BioAnalyzer, 500 ng of total RNA with RIN values of 10 underwent polyA selection and library preparation using the TruSeq Stranded mRNA LT Kit (Illumina catalog # RS-122-2102) according to instructions provided by the manufacturer with 8 cycles of PCR. Samples were barcoded and run on a NovaSeq 6000 in a PE100 run, using the NovaSeq 6000 S4 Reagent Kit (200 Cycles) (Illumina). An average of 204 million paired reads was generated per sample. Ribosomal reads represented 0.09-0.32% of the total reads generated and the percent of mRNA bases averaged 92%. Bulk RNA-seq samples were sequenced using the Illumina Novaseq 6000 platform, generating 100-base pair paired-end reads. FASTQ files were quality-checked using FastQC v0.12.0 and trimmed with Trim Galore v0.6.10 to remove adapter sequences and read ends with Phred scores below 15. STAR v2.7.0a was used to align the FASTQ files to the hg38 reference genome. The resulting SAM files were sorted using the sort function from Samtools v1.19.2 and indexed using the index function to generate BAM index files.Gene expression was quantified using featureCounts v2.0.6 (Subread package), configured to count fragments assigned to exon features, and summarized by gene name based on the GENCODE v45 gene annotations. Differential expression analysis was performed using DESeq2 v1.42.0.

To evaluate the ISR gene signature, the ISR gene signature scores were calculated from the normalized count matrix of the bulk RNA-seq data with the UCell algorithm in R. This algorithm calculates gene enrichment scores for each replicate of the Ctrl or MGO-treated cells. The ISR gene signature is defined by the curated list of ISR genes.^55^ Only genes with a status ‘Upregulated’ in IRS gene list (n=224 + ATF4) were included in the analyses.

### RT-PCR

WT HEK293T cells were cultured to 80% confluency in a 6 well plate. Media was supplemented with 0 or 200 nM ISRIB for 30 minutes prior to transfection with 5 µg of mRNA. 4 hours post-transfection, cells were washed once with warm PBS and collected. RNA was purified using the RNeasy Mini Kit (Qiagen catalog #74104) according to manufacturer instructions. mRNA was reverse-transcribed using the High-Capacity RNA-to-cDNA Kit (Applied Biosystems Catalog #4387406) according to manufacturer instructions. RNA and cDNA levels were quantified using a NanoDrop™ One/OneC Microvolume UV Vis Spectrophotometer (Thermo Scientific).

Quantitative PCR reactions were performed using the iTaq Universal SYBR green supermix (Biorad Catalog #172-5121) with each 10 uL reaction containing 100 ng cDNA and 500 nM of both forward and reverse primers for the reaction. The amplification program was initiated at 95 °C for 3 minutes followed by 40 cycles 95 °C for 10 seconds and 60 °C for 30 seconds. Relative RNA abundances were deduced from ΔΔC_T values_, normalized to HPRT1 mRNA abundance, and compared to its respective control sample.

### CELL CYCLE ANALYSIS

WT HEK293T cells were seeded at 80% in a 96-well plate and treated with the indicated concentration of MGO for 1 hour. At endpoint, cells were washed with PBS to remove any residual MGO. Viability staining was done using LIVE/DEAD Violet following the user instructions. Cells were then fixed using the FoxP3 Transcription Factor Staining kit and stained with DAPI, to determine cell cycle, for 30 minutes. Measurement of DNA content was determined by flow analysis on the Cytek Aurora. Cell cycle distribution analyses was conducted using FlowJo.

### GLUCOSE-STIMULATED INSULIN SECRETION

MIN6 cells were treated with MGO for 1 before incubation in Krebs–Ringer containing 2.8 mM D-glucose for 2 h. Glucose-stimulated insulin secretion was assessed by incubating cells in Krebs–Ringer buffer containing either 2.8 mM (low glucose) or 16.7 mM (high glucose) D-glucose for 1 h. Following stimulation, supernatants were collected for insulin measurement. Cells were subsequently lysed, and total protein content was determined using a BSA assay. Insulin levels were quantified using an ELISA kit (ALPCO, Salem, NH). Glucose-stimulated insulin secretion (GSIS) was calculated as fold change relative to untreated control conditions. For mouse studies, insulin levels were measured from 10 µL of serum using an ELISA kit (ALPCO, Salem, NH), according to the manufacturer’s instructions.

#### Quantification and statistical analysis

Statistical analyses were performed in GraphPad Prism v10.6.0. Specific tests used, error bar information, and number of experimental repeats can be found in figure legends. Data are displayed as individual values and/or mean values with error bars signifying the standard deviation (SD) No statistical methods were used to predetermine sample sizes, and investigators were not blinded during experiments or analysis.

## Acknowledgments

We thank members of the David lab and Heeseon An for helpful discussions. MIN6 cells were a kind gift from Danwei Huangfu Lab (MSKCC) and we thank Tamara Casteels for the help with the MIN6 cells and insulin secretion assays. Wild-type U2OS and the ΔZAK U2OS cells were a kind gift from Simon Bekker-Jensen Lab (University of Copenhagen) and we thank Melanie Blasius for the help with the USOS cells. We acknowledge the use of the Memorial Sloan Kettering Cancer Center Integrated Genomics Operation Core and the Memorial Sloan Kettering Cancer Center Molecular Cytology, funded by the NCI Cancer Center Support Grant (CCSG, P30 CA08748). Work in the Y.D. lab is supported by the Josie Robertson Foundation, the Pershing Square Sohn Cancer Research Alliance, the NIH (CCSG core grant P30 CA008748, MSK SPORE P50 CA197, and R21 DA044767), the Parker Institute for Cancer Immunotherapy (PICI) and the Anna Fuller Trust. In addition, the Y.D. lab is supported by Mr. William H. Goodwin and Mrs. Alice Goodwin, the Commonwealth Foundation for Cancer Research, and the Center for Experimental Therapeutics at Memorial Sloan Kettering Cancer Center. A.K. is supported by the Swiss National Science Foundation through the SNF Postdoc Mobility Fellowship (214384) and the Basic Research Innovation Scholars Fellowship by the Starr Foundation.

## Author Contributions

A.K., K.N., and Y.D. conceived the study and designed experiments. A.K. and K.N. performed experiments, analyzed data, and prepared figures. N.K. and J.G. performed LCMS analyses. M.D., F.F., J.R., and O.A.-Q. performed bioinformatic analyses. N.C., R.K., and V.R.S. performed mouse experiments. S.M. performed *in vitro* ribosome collision assays. N.W. performed cell-cycle arrest assays. X. M. performed ZAK-phosphorylation assays. A.D. contributed to Keth-seq analyses. C.E., M.G.K., A.S. and K.S. helped with experiments related to ribosome stalling. Y.X. assisted with experiments. A.K., K.N., and Y.D. wrote the manuscript. All authors reviewed and approved the final manuscript.

