## Supplementary material for "Non-enzymatic RNA Glycation is a Metabolic Sensor of Cellular Stress": SI_Figures

**Affiliations:**

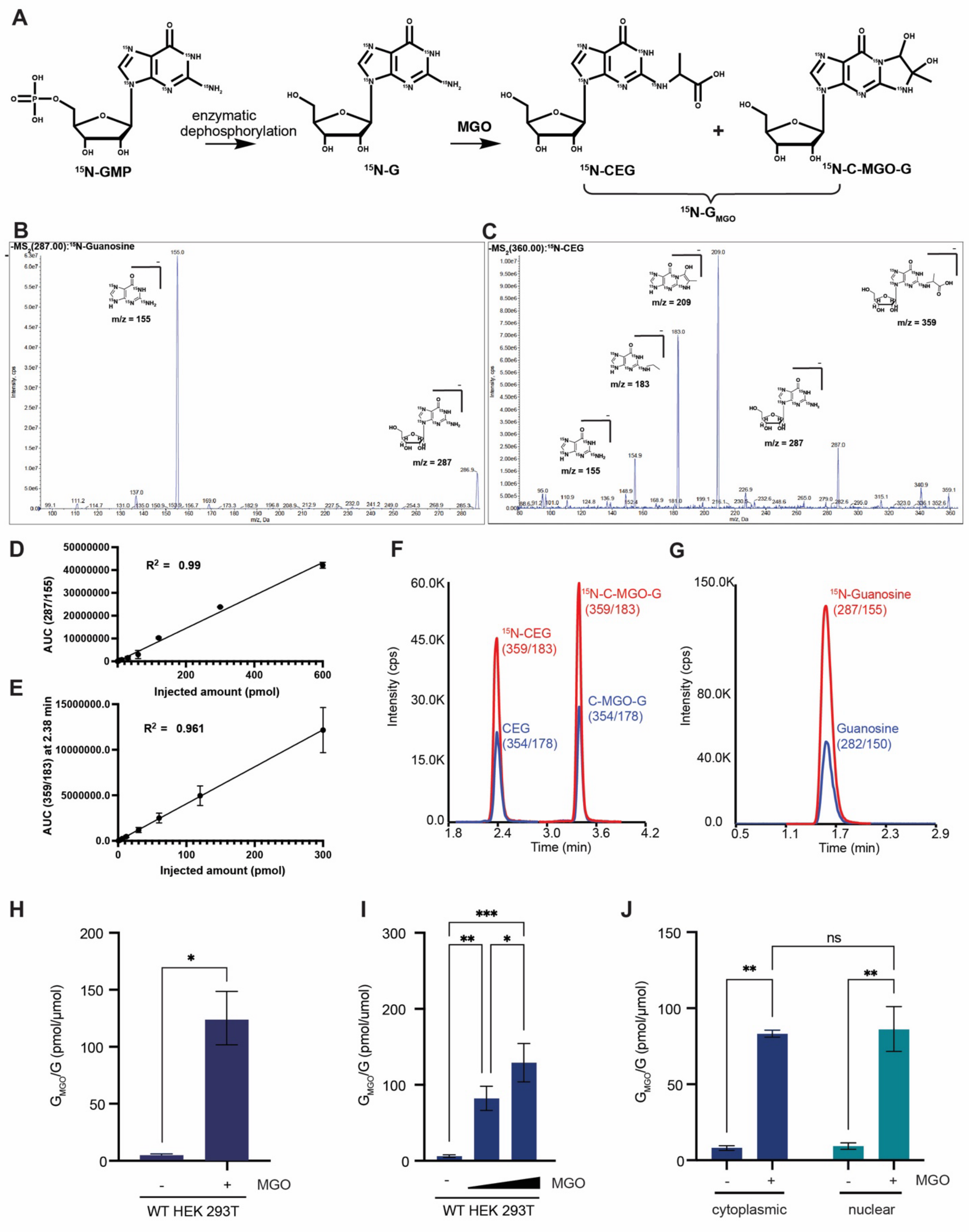

A
15N-GMP
enzymatic dephosphorylation
15N-G
MGO
15N-CEG
15N-C-MGO-G
15N-GMGO
B
-MS2(287.00):15N-Guanosine
m/z = 155
m/z = 287
C
-MS2(360.00):15N-CEG
m/z = 209
m/z = 359
m/z = 183
m/z = 287
m/z = 155
D
R2 = 0.99
AUC (287/155)
Injected amount (pmol)
E
R2 = 0.961
AUC (359/183) at 2.38 min
Injected amount (pmol)
F
15N-CEG (359/183)
15N-C-MGO-G (359/183)
CEG (354/178)
C-MGO-G (354/178)
Intensity (cps)
Time (min)
G
15N-Guanosine (287/155)
Guanosine (282/150)
Intensity (cps)
Time (min)
H
GMGO/G (pmol/µmol)
MGO
WT HEK 293T
I
GMGO/G (pmol/umol)
MGO
WT HEK 293T
J
GMGO/G (pmol/µmol)
ns
MGO
cytoplasmic
nuclear

**<u>Fig. S1</u>: Optimizing LCMS protocol to detect and quantify RNA glycation (related to Figure 1).** (A) Synthesis route for isotopically labelled standards $^{15}$N-G and $^{15}$N-$G_{MGO}$ (B) MS2 spectra of 10 μM $^{15}$N-G (C) MS2 spectra of 10 μM $^{15}$N-$G_{MGO}$ (D) Linear regression analysis of injected $^{15}$N-G (pmol) and Area Under Curve (AUC) of 287/155. (E) Linear regression analysis of injected $^{15}$N-$G_{MGO}$ (pmol) and Area Under Curve (AUC) of 359/183 at 2.38 mins (CEG). (F) Overlay of LC-MS chromatograph of $^{15}$N-$G_{MGO}$ and $G_{MGO}$ (G) Overlay of LC-MS chromatograph of $^{15}$N-G and G. (H) $G_{MGO}$ levels of total RNA isolated from WT HEK293T cells treated with 0 and 120 μM MGO for 1 hour. Quantified by LCMS to isotopically labeled standards (n=2). Statistical significance was assessed using a two-tailed unpaired t-test (*$P$< 0.0332; **$P$< 0.0021; ***$P$< 0.0002; ****$P$< 0.0001). (I) $G_{MGO}$ levels of total RNA isolated from WT HEK293T cells treated with 0, 60 μM and 120 μM MGO for 4 hours. Quantified by LCMS to isotopically labeled standards (n=3). Statistical significance was assessed using a Two-way ANOVA with Tukey's multiple comparison test (*$P$< 0.0332; **$P$< 0.0021; ***$P$< 0.0002; ****$P$< 0.0001). All data are presented as mean ± SD. (J) $G_{MGO}$ levels of cytoplasmic and nuclear enriched RNA isolated from ΔGLO1 HEK293T cells treated with 0 or 60 μM MGO for 4 hours. Quantified by LCMS to isotopically labeled standards (n=3). Statistical significance was assessed using a Two-way ANOVA with Tukey's multiple comparison test (*$P$< 0.0332; **$P$< 0.0021; ***$P$< 0.0002; ****$P$< 0.0001).

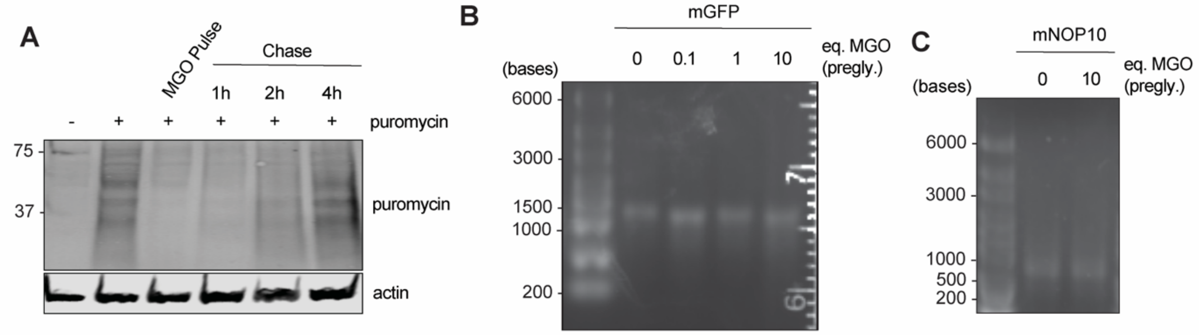


**Fig. S2: RNA glycation stalls translation (related to Figure 3).** (A) Immunoblot analysis of puromycin incorporation in WT HEK293T cells following an MGO pulse–chase experiment. Cells were pulsed with 0.47 µM MGO for 1 hour, washed, and incubated in fresh medium for the indicated chase times. Puromycin (5 µg/mL final concentration) was added during the last 15 min of each time point to label nascent polypeptides (n=3). (B) Native RNA gel of mGFP incubated with indicated amounts of MGO for 16h at 37 °C and purified. (C) Native RNA gel for mNOP10 incubated with indicated amounts of MGO for 16h at 37 °C and purified.

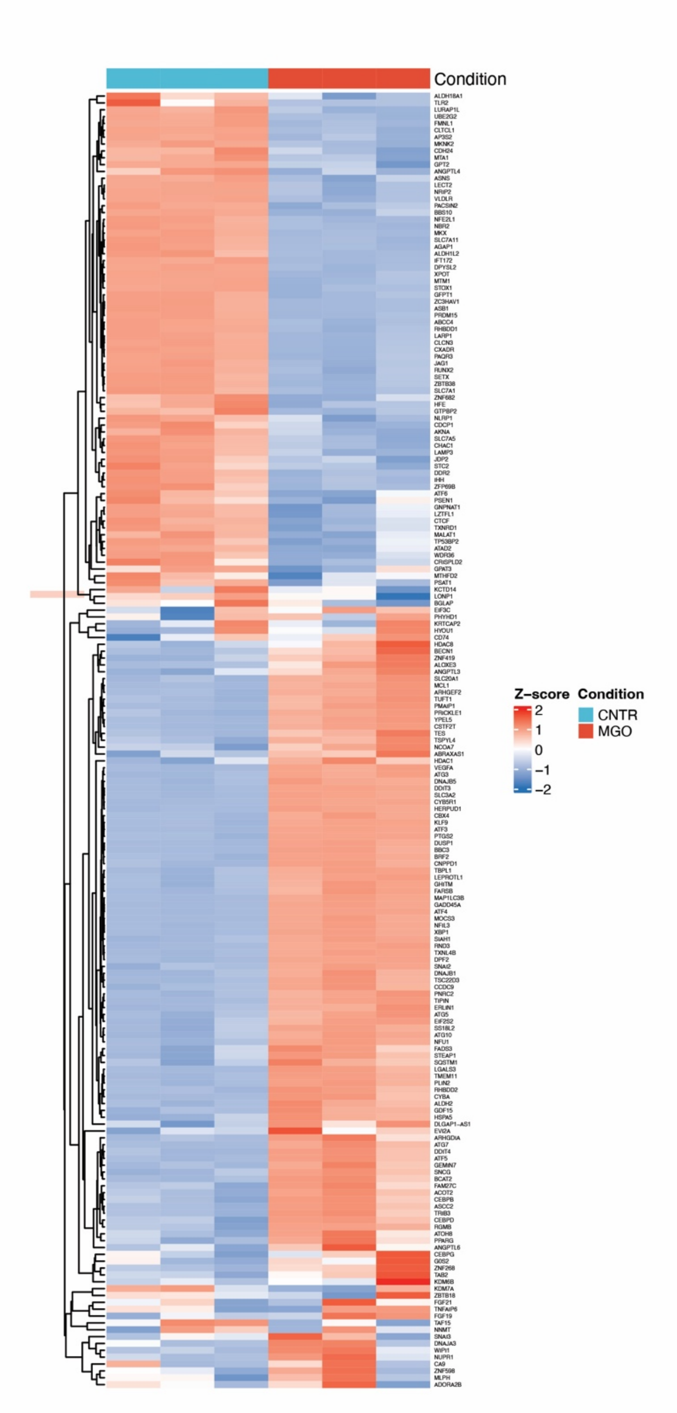


**<u>Fig. S3</u>: RNA glycation induces ribosome collision and activation of the integrated stress response (related to Figure 4).** Integrated stress response (ISR) gene expression signature derived from RNA-seq analysis of control and MGO-treated ΔGLO1 HEK293T cells (60 μM MGO, 4 h), highlighting ISR-dependent transcripts.

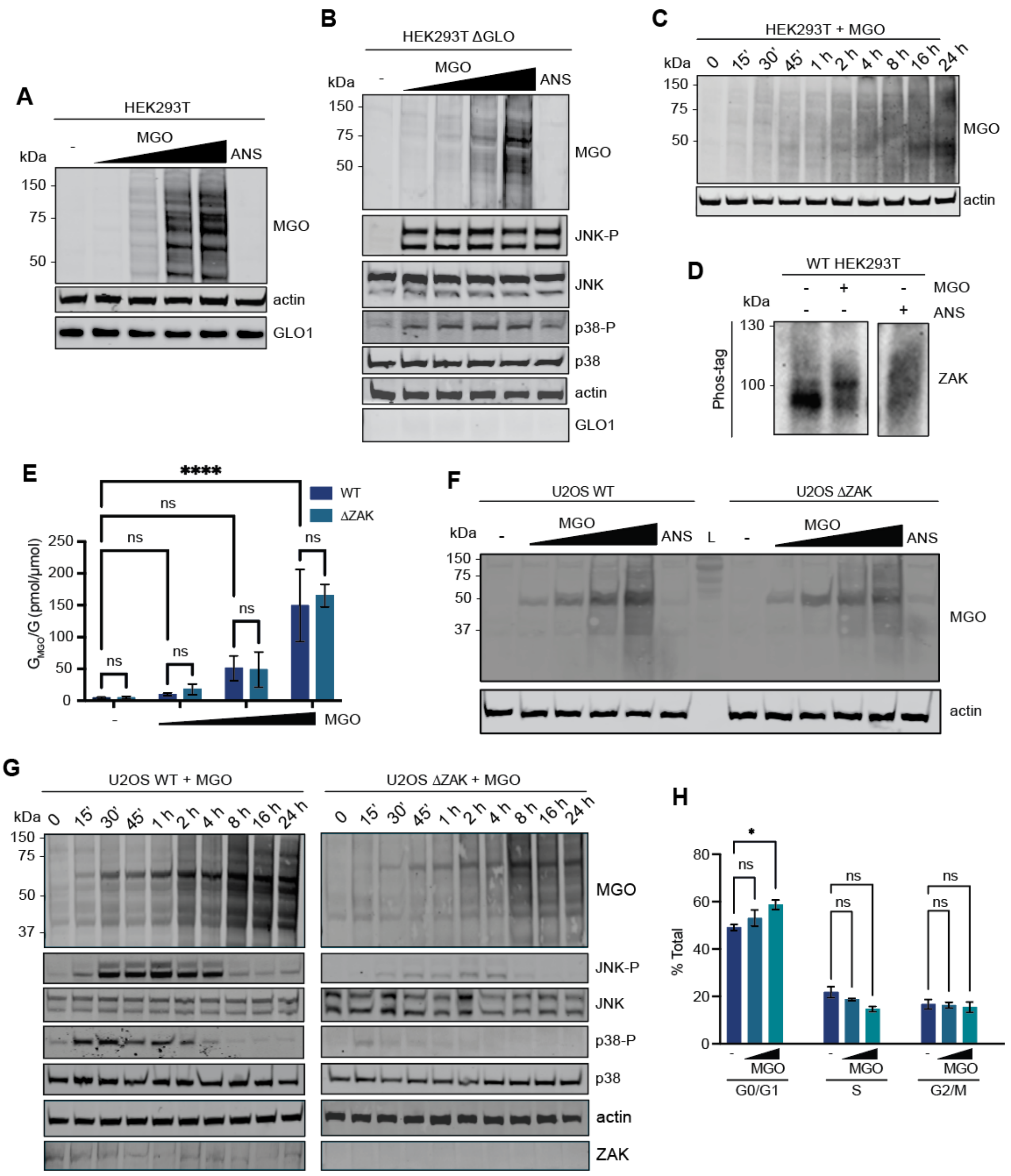


**Fig. S4: RNA glycation activates the ribotoxic stress response (related to Figure 5).** (A) Immunoblot analysis of WT HEK 293T cells treated for 1h with 0-0.35 mM MGO or 1 µg/mL anisomycin (positive control for RSR activation) (n=3). (B) Immunoblot analysis of ΔGLO1 HEK 293T cells treated for 1h with 0-0.12 mM MGO or 1 µg/mL anisomycin (positive control for RSR activation) (n=3). (C) Immunoblot analysis of WT HEK 293T cells after treatment with 0.35 mM MGO for the indicated time. (D) Low contrast image of immunoblot of Phostag gel for ZAK phosphorylation in WT HEK293T cells treated with or without 0.24 mM MGO for 1 h, or with 1 µg/mL anisomycin for 1 h (n=3). (E) $G_{MGO}$ levels of total RNA isolated from WT and ΔZAK U2OS cells treated with 0-240 µM MGO for 1 hour. Quantified by LCMS to isotopically labeled standards (n=3). Statistical significance was assessed using a two-way ANOVA with Tukey's multiple comparison test (*$P$< 0.0332; **$P$< 0.0021; ***$P$< 0.0002; ****$P$< 0.0001). (F) Immunoblot analysis of WT and ΔZAK U2OS cells treated for 1 hour with 0-0.24 mM MGO or 1 µg/mL anisomycin (n=3). (G) Immunoblot analysis of WT (left panel) and ΔZAK (right panel) U2OS cells after treatment with 0.35 mM MGO for the indicated time (H) Quantitative analysis of the percentage of WT HEK293T cells in each phase after treatment with 0-0.24 mM MGO for 1 hour. (n=3). Statistical significance was assessed using a Two-way ANOVA with Tukey's multiple comparison test (*$P$< 0.0332; **$P$< 0.0021; ***$P$< 0.0002; ****$P$< 0.0001).

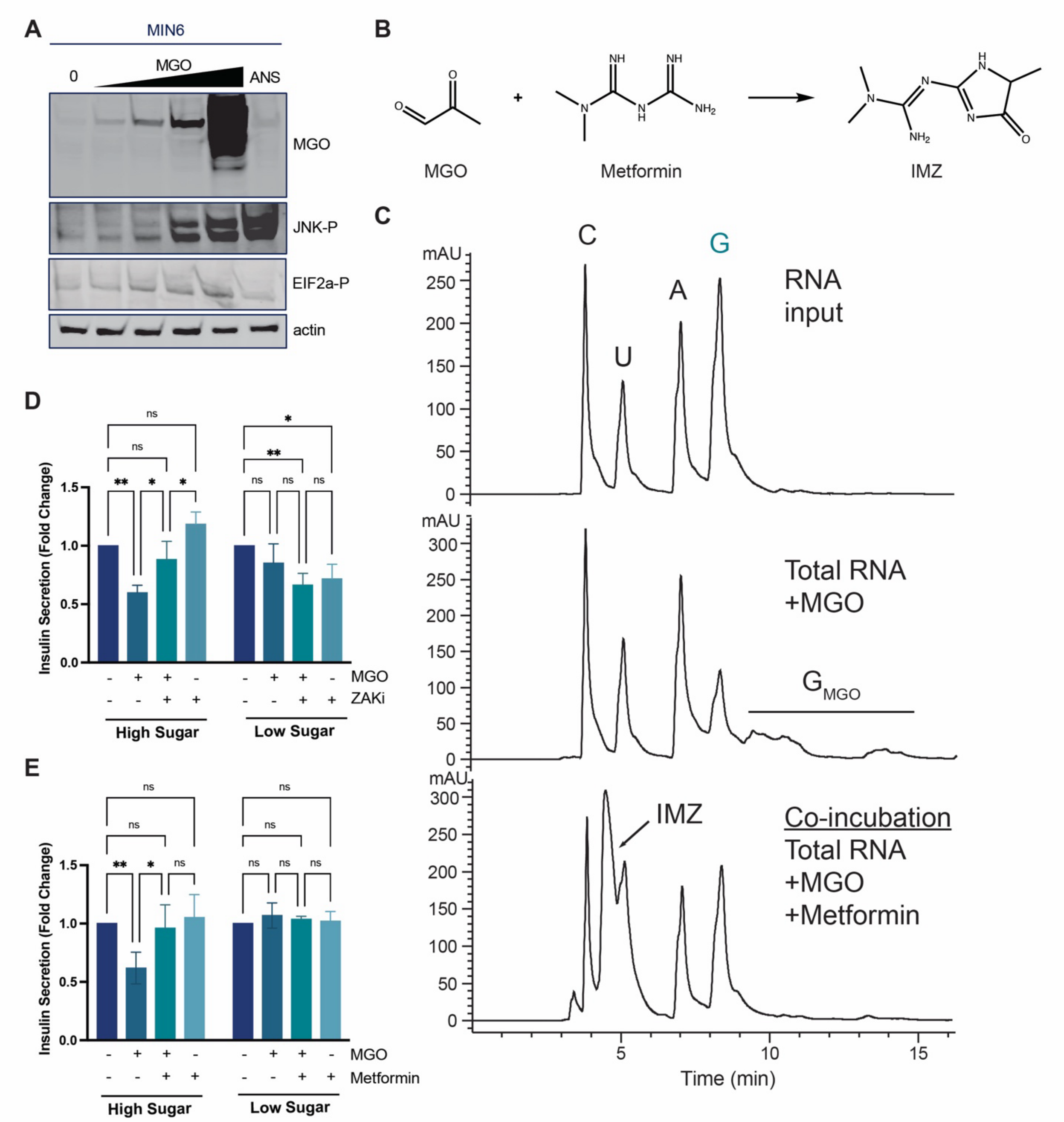


**Fig. S5: MGO induces the RSR and ISR in β-cells to regulate insulin secretion (related to Figure 6).** (A) Immunoblot analysis of MIN6 cells treated for 1h with 0-0.24 mM MGO or 1 μg/mL anisomycin (positive control for RSR activation) (n=3). (B) Reaction scheme of MGO and metformin. (D) Glucose-stimulated insulin secretion (GSIS) in MIN6 cells under high glucose (16.7 mM) stimulation conditions. Cells were preincubated with 0 or 0.24 mM MGO for 1 h, with or without ZAKi pretreatment (1 μM, 30 min). Data are shown as fold change relative to untreated controls (n=3). Statistical significance was assessed using a Two-way ANOVA with Tukey's multiple comparison test (**P*< 0.0332; ***P*< 0.0021; ****P*<

0.0002; ****$P$< 0.0001). (E) GSIS in MIN6 cells treated with 0.24 mM MGO for 1h in the presence or absence of metformin pretreatment (1 mM for 1 h), (n=3), Statistical significance was assessed using a Two-way ANOVA with Tukey's multiple comparison test (*$P$< 0.0332; **$P$< 0.0021; ***$P$< 0.0002; ****$P$< 0.0001).
